# High response diversity and conspecific density-dependence, not species interactions, drive dynamics of coral reef fish communities

**DOI:** 10.1101/2024.01.23.576565

**Authors:** Alfonso Ruiz-Moreno, Michael J. Emslie, Sean R. Connolly

**Affiliations:** College of Science and Engineering, James Cook University, Townsville, QLD 4811 Australia; Australian Institute of Marine Science, PMB 3 Townsville MC, Townsville, QLD 4810, Australia; Smithsonian Tropical Research Institute, Panama City, Panama

**Keywords:** Species interactions, response diversity, reef fish, latent variables, regularized horseshoe, community dynamics, time series

## Abstract

Species-to-species and species-to-environment interactions are key drivers of community dynamics. Disentangling these drivers in species-rich assemblages is challenging due to the high number of potentially interacting species (the “curse of dimensionality”). We develop a process-based model that quantifies how intraspecific and interspecific interactions, and species’ covarying responses to environmental fluctuations, jointly drive community dynamics. We fit the model to reef fish abundance time series from 41 reefs of Australia’s Great Barrier Reef. We found that fluctuating relative abundances are driven by species’ heterogenous responses to environmental fluctuations, whereas interspecific interactions are negligible. Species differences in long-term average abundances are driven by interspecific variation in the magnitudes of both conspecific density-dependence and density-independent growth rates. This study introduces a novel approach to overcoming the curse of dimensionality, which reveals highly individualistic dynamics in coral reef fish communities that imply a high level of niche structure.

## Introduction

Understanding what drives variation in abundances of species in space and time is a core aim of ecology (Krebs 2009). Populations respond to a combination of density-dependent and density-independent processes (Ohlberger *et al*. 2014; Sæther *et al*. 2016; Thibaut & Connolly 2020). Interspecific interactions, such as competition and facilitation, also can influence species-abundance dynamics (Butterfield 2009; Roughgarden 1974; Tilman 1994), and have been hypothesized to be particularly important in the tropics (Dobzhansky 1950; Schemske *et al*. 2009; Paquette & Hargreaves 2021). Species-to-environment interactions, such as variable species responses to environmental fluctuations, and covariations in those responses, also drive relative abundance dynamics. (Elmqvist *et al*. 2003; Thibaut *et al*. 2012; Thibaut & Connolly 2013). Particularly, if species are not perfectly positively correlated in their responses to environmental fluctuations, their fluctuations in abundance will tend to be less pronounced at the community level than at the individual species level (a phenomenon referred as “response diversity”, Elmqvist *et al*. 2003).

Evaluating effects of interspecific and species-to-environment interactions on community dynamics is particularly challenging in species-rich communities, due to the “curse of dimensionality”. That is, the number of potential interspecific interaction strengths, and covariances in species’ response to environmental fluctuations, that need to be estimated, increases quadratically with the number of species in the community (e.g., from 12 to 90 to 380 interaction terms and 6 to 45 to 190 environmental covariances as a community increases from 4 to 10 to 20 species).

In the absence of practical methods to estimate so many parameters, either experimentally or via time-series analysis, theoretical community ecologists have developed highly parsimonious biodiversity models that make strong simplifying assumptions about community dynamics. At the extreme, neutral theory of biodiversity assumes that individuals are identical regardless of species identity, and all the variability in a community is driven by demographic stochasticity — random variation in the fates of individuals (e.g., birth, death and dispersal events) (Hubbell 2001). However, neutral models’ ability to explain biodiversity patterns in real communities has been challenged (e.g., Brown *et al*. 2013; Chisholm *et al*. 2014; Connolly *et al*. 2014).

Somewhat less restrictive, the stochastic community-dynamic theory of Engen and colleagues (Engen & Lande 1996; Engen *et al*. 2002; hereafter the “Engen model”) allows for species differences in demographic rates and their fluctuations with environmental conditions. However, this theory still has restrictive assumptions. Specifically, the temporal variance of fluctuations in the density independent population growth rate, and the strength of intraspecific density dependence are assumed to be equal for all species (*contra*, e.g., Comita *et al*. 2010; Johnson *et al*. 2012; LaManna *et al*. 2017; Mangan *et al*. 2010). Additionally, interspecific interactions are assumed to be negligible and species’ responses to environmental fluctuations are independent (Engen & Lande 1996), or at least equally positively correlated (Appendix B in Engen *et al*. 2002). How strongly these assumptions are violated, and how robust the inferences made from such models (e.g., Engen *et al*. 2002; Solbu *et al*. 2018), has been assessed only for a narrow range of parameter values (Tsai *et al*. 2022).

These two dimension-reduction approaches have been applied previously to coral reef fish assemblages. Static analyses of patterns of commonness and rarity reveal these communities do not follow species-abundance distributions expected from neutral dynamics (Connolly *et al*. 2014; Connolly *et al*. 2017). Furthermore, analysis of temporal dynamics of species-abundance distributions, using the Engen model, suggests that most of the variability in coral fish abundances is due to persistent heterogeneity in demographic characteristics among species, with smaller contribution due to environmental fluctuations (Tsai *et al*. 2022). However, species differences in the strength of density-dependence has been hypothesized to be an important driver of variation in abundance in other high-diversity assemblages like tropical forests (Comita *et al*. 2010; Johnson *et al*. 2012; LaManna *et al*. 2017; Mangan et al. 2010), and differential sensitivity of species to environmental fluctuations have been widely documented, including for reef fishes (Emslie *et al*. 2011; Hoey *et al*. 2016; Pratchett *et al*. 2011, 2015). Similarly, experimental studies have suggested potentially strong interactions among some reef fishes (Ebersole 1977; Jones 2005; Robertson 1996; Shulman 1985), contrary to the assumption of negligible interspecific interactions. Ideally, to draw robust inferences about community structure, we would like to confront reef fish community data with models that can account for such heterogeneities and interactions, where they are present.

Some promising approaches allowing greater interspecific heterogeneity in models for high-diversity communities have been developed. For example, joint species distribution models use observed environmental and latent predictors to model species’ fluctuations in abundances as functions of a smaller number of predictors, rather than trying to independently estimate pairwise interaction terms or environmental covariances (Hui *et al*. 2015; Ovaskainen *et al*. 2017a; Warton *et al*. 2015). While this can identify the contribution of specific environmental predictors, the latent predictors could represent either species interactions or the effects of unmeasured environmental drivers (see methods for a more detailed example of the latent variable approach) (Ovaskainen *et al*. 2017a). Additionally, such models can only detect symmetric species associations (Hui *et al*. 2015; Ovaskainen *et al*. 2017a; Warton *et al*. 2015), which makes sense in the context of covariances in responses to environmental fluctuations, but not species interactions (e.g., predator-prey interactions, where one species benefits while the other is harmed).

This study aims to evaluate the importance of among-species heterogeneity in demographic rates (particularly the strength of density dependence, and the sensitivity of species’ density-independent growth rates to environmental fluctuations), species interactions, and response diversity as drivers of the temporal dynamics of reef fish assemblages on the Great Barrier Reef (GBR), Australia. Specifically, we develop a mechanistic model that is tractable, but can estimate both species interactions and the variances and covariances of species’ response to environmental fluctuations, without strong homogeneity assumptions, while making plausible biological assumptions about how those heterogeneous quantities are distributed among species. We fit this model to reef fish assemblage data, then evaluate the magnitude and importance of species interactions and response diversity as drivers of changes in abundance, and the relative importance of heterogeneity in density-independent and density-dependent demographic parameters in driving persistent variation in abundances among species. We also test our approach using simulated data, to evaluate whether it can successfully recover the parameters used to simulate the data, and in particular, to distinguish between covariation in species’ abundances that is mediated by species interactions versus environmental fluctuations. Our findings highlight highly heterogeneous and individualistic dynamics, with species interactions overwhelmingly negligibly small and response diversity relatively high. We also find that the substantial heterogeneity in species’ long-term abundances is driven approximately equally by interspecific differences in density-dependent and density-independent components of population growth.

## Methods

### Data collection

The GBR’s reef fish communities have been surveyed by the Australian Institute of Marine Science’ Long Term Monitoring Program (LTMP) since 1995 (Emslie *et al*. 2020). Underwater visual surveys were conducted annually on the same 41 reefs (Fig. 1) between 1995 to 2005, so here we focus on these reefs for this 11-year period. At each reef, there were three sites on the reef slope, usually on the north-east flank of the reef. Each site contained five permanently marked 50m transects, approximately parallel to the reef crest between 6m and 9m. Observers recorded abundances of 208 species of reef fishes from 9 families: *Labridae* (including *Scarine* parrotfishes), *Pomacentridae*, *Siganidae*, *Chaetodontidae*, *Acanthuridae*, *Serranidae*, *Lutjanidae*, *Lenthrinidae* and *Zanclidae*. Pomacentrids were counted on 50m-by-1m transects and all other families on 50m-by-5m transects (see Emslie & Cheal 2018 for detailed methodology).

**Figure 1.**
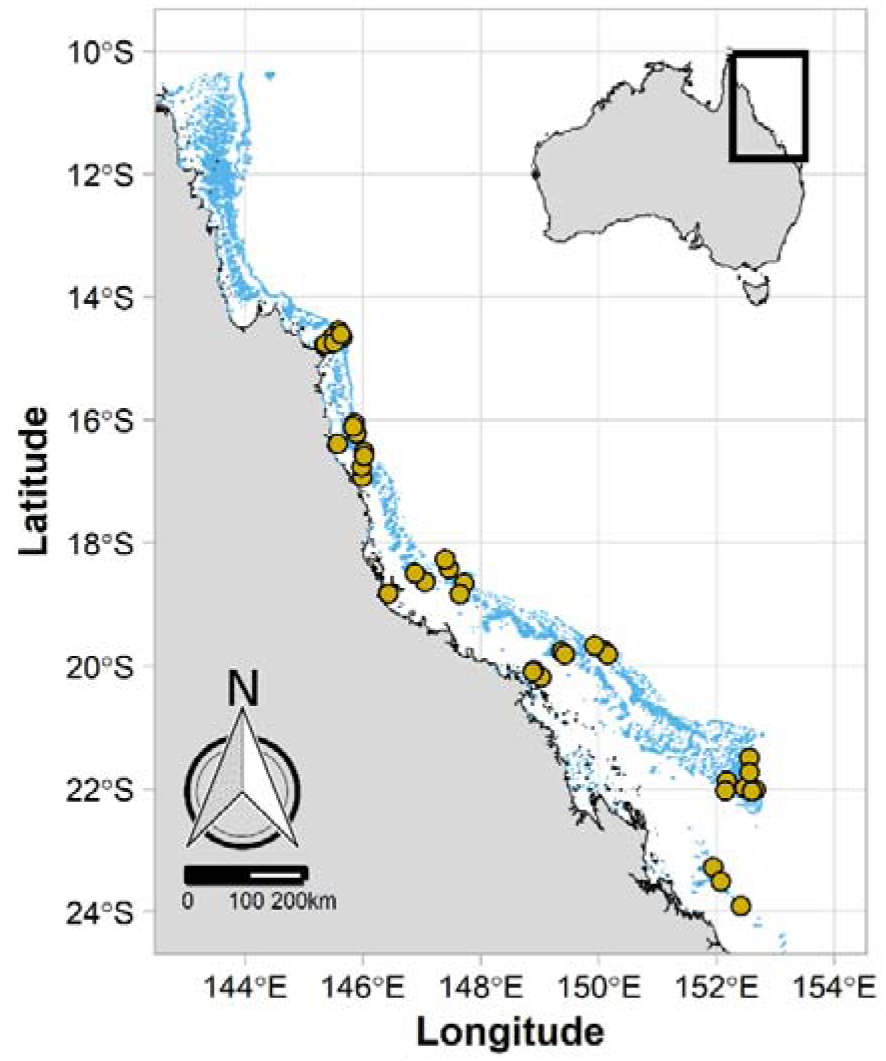
Map of the GBR showing the 41 sampled coral reefs in our analysis as yellow points. Mainland Australia and islands are represented in grey and coral reefs and cays are in light blue.

### The model

The dynamics of abundances are assumed to follow the multivariate Gompertz model (Ives *et al*. 2003), which has been used previously to model the dynamics of reef fishes (Thibaut *et al*. 2012; Tsai *et al*. 2022), and which characterizes the density-dependent dynamics of reef fishes better than models of logistic form, such as the Lotka-Volterra model (Thibaut *et al*. 2012). The Gompertz model follows:

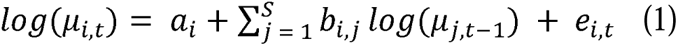

or, in matrix form:

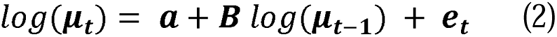

where log(μ**_t_**) is a vector containing species’ estimated log-abundances at time *t,* μ**_t_** *= (*μ*_1,t_,* μ*_2,t_,* μ*_3,t_,…,* μ*_i,t_)*. ***a*** is a vector containing species’ estimated intrinsic rates of increase, ***a*** = (*a_1_*, *a_2_*, *a_3_*,…, *a_i_*). **B** is a species-by-species interaction matrix whose off-diagonal elements, *b_ij_*, indicate the effect of the abundance of species *j* on the per capita population growth rate of species *i* (*b_ij_* = 0 for no interaction; *b_ij_* < 0 for negative effects (e.g. competition); *b_ij_* > 0 for positive effects (e.g. facilitation)), and whose diagonal elements, *b_ii_*, represent the effect of the abundance of species *i* on its own population growth (*b_ii_* = 1 for density-independent growth; 0 < *b_ii_*< 1 implies compensatory density-dependence, *b_ii_* < 0 implies over-compensatory density-dependence). **e_t_** is a vector of process error for each species, **e_t_** = (*e_1,t_, e_2,t_, e_3,t_,…, e_i,t_*), which has a multivariate normal distribution with mean vector **0** and covariance matrix Σ. This represents stochastic fluctuations in the intrinsic growth rate from year-to-year.

We modelled observed fish counts as Poisson-distributed:

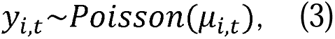

where *y_i,t_*is the observed count of fish of species *i* at time *t,* μ*_i,t_* is the (unobserved) abundance of species *i* at time *t* from eq. 1.

Due to the use of different transect sizes to count Pomacentrids (50m x 1m transect) and non-Pomacentrids (50m x 5m transect), we modelled the Pomacentrid counts as:

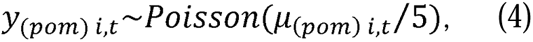

where *y_(pom)i,t_*is the observed number of fish of the pomacentrid species *i* at time *t,* μ*_(pom)i,t_* is the mean abundance of the pomacentrid species *i* at time *t* per 250m2 (the area of the larger transect). The division by 5 accounts for the fact that pomacentrids were counted on transects that were a fifth the size of the normal transects. This obviates the need to exclude information by subsampling fishes counted on the larger transects (e.g., as in Connolly *et al*. 2005, 2009, 2017; Tsai *et al*. 2022).

### Reducing Dimensionality of the Model

As species richness increases, the number of parameters in the interaction matrix **B** and the covariance matrix Σ increase quadratically. The interaction matrix **B** has *S^2^* free parameters, where S is the number of species. The covariance matrix Σ has *S(S+1)/2* free parameters. Estimating that many free parameters in speciose assemblages would require very long time-series data, which generally do not exist for community time series, particularly in marine systems.

To reduce the dimensionality of the covariance matrix, Σ, we used a factor analysis approach, similar to two previous analyses of high-diversity time series (Ovaskaninen *et al*. 2017b; Sandal *et al*. 2022). This approach assumes that the observed data can be explained by a small number of latent variables, *D (D << S)*, while still explaining the covariance of the observed data (Hui *et al*. 2015; Ovaskainen *et al*. 2017a; Warton *et al*. 2015). The logic is that environmentally-mediated fluctuations in abundances should be driven by a common set of environmental drivers, which in general may not be known or measured. Therefore, the variance-covariance matrix was estimated as

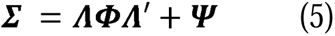

where Λ is an *S x D* matrix of factor loadings, which can be interpreted as the response of species to the unknown drivers (i.e., latent variables). Φ is the variance-covariance matrix of the latent variables and it is a *D*-by-D matrix (see Appendix S1). The covariance matrix Ψ is a diagonal matrix explaining the remaining variation (residual error) not captured by the factor loadings and the latent factors. The correlation in species responses to environmental fluctuations, **P**, can be calculated from the variance-covariance Σ as 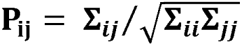. This changes the number of free parameters in the covariance matrix (**Σ = ΛΦΛ’+Ψ)** from *S(S+1)/2* to *D(S+(1-D)/2)*. Provided that *D<<S*, this substantially reduces the number of parameters required to calibrate the covariance matrix Σ (See Appendix S1 for further details and Fig. 2 for prior choices and model structure).

**Figure 2.**
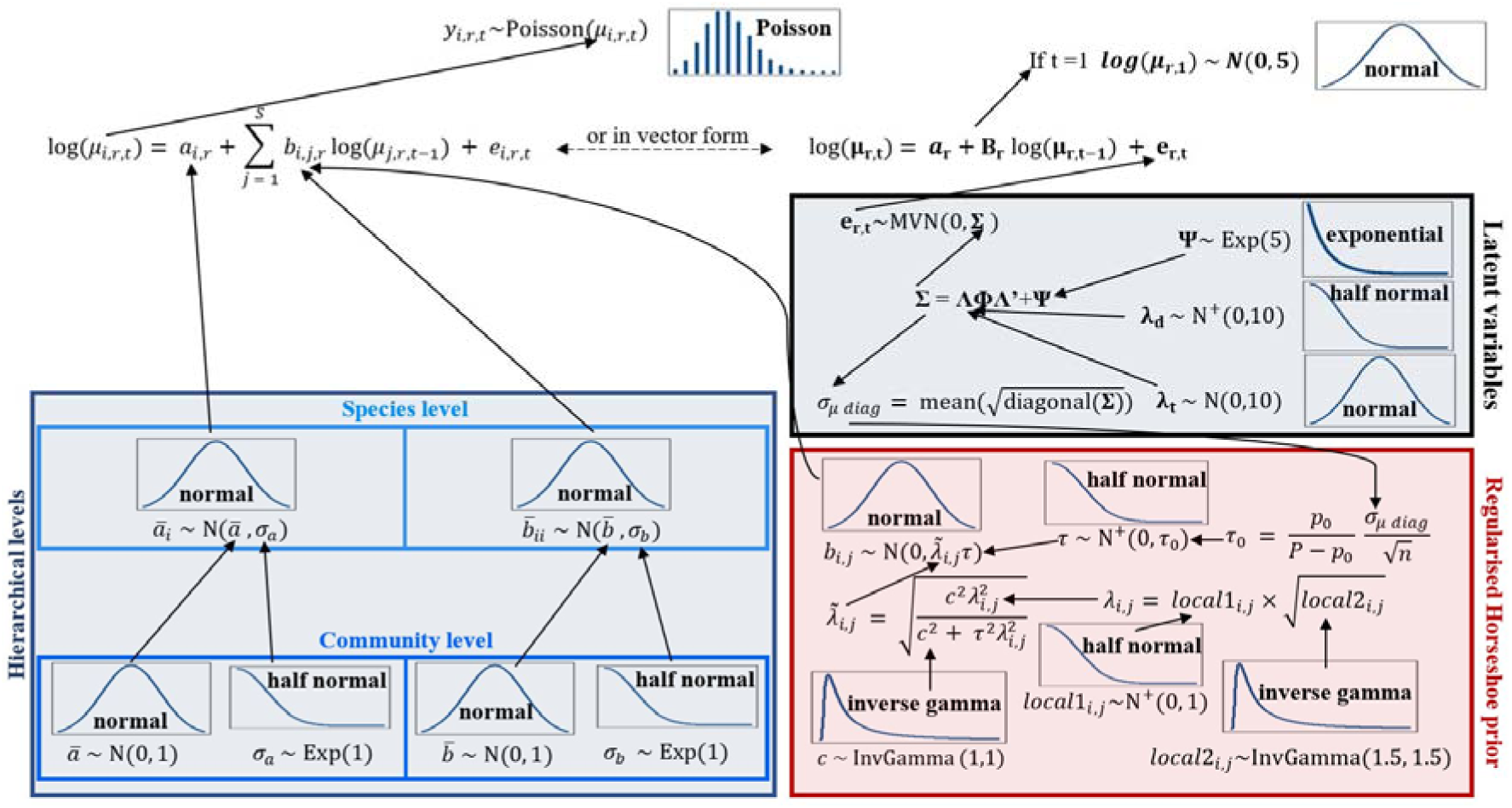
Schematic of the Poisson multivariate autoregressive Gompertz model. The blue box shows the random effects for the intrinsic growth rate and the within-species density dependence. There are two levels: the metacommunity parameters, and the parameters for each species. The red box shows the regularised horseshoe prior used to estimate the off-diagonal elements of the interaction matrix (between species density dependence). The black box shows the factor analysis component used to estimate the variance-covariance matrix. Note that λ_d_ stands for the diagonal elements of the factor loadings matrix Λ, whereas λ_t_ stands for the lower triangular elements of the factor loadings matrix Λ. Normal distributions are denoted with the standard deviation formulation (e.g., N(0, 2) indicates a normal distribution with mean 0 and standard deviation of 2, not a variance of 2).

With respect to the interaction matrix, **B**, different studies have found evidence of weak or negligible interspecific interactions versus strong competitive intraspecific interactions in time series data for marine microorganism, moth, temperate fish, crustacean, bird and rodent communities (Mutshinda *et al*. 2009; Ovaskaninen *et al*. 2017b; Sandal *et al*. 2022; but see Almaraz & Oro 2011). In contrast, some of community ecology’s most classic studies identify species interactions with important demographic consequences (e.g., competitive [Connell 1961], keystone species [Paine 1966], predator-prey cycles [Stenseth *et al*. 1997; Krebs *et al*. 2017]), and some studies suggest potentially strong effects of reef fishes on one another (Ebersole 1977; Jones 2005; Robertson 1996; Shulman 1985).

Therefore, we seek an approach in which most interactions will be weak or negligible, but which allows some interactions to be strong, and potentially asymmetric. To do this, we implemented the “regularised horseshoe prior” (Piironen & Vehtari 2017) as a prior distribution for our interspecific interaction terms. This distribution has high density around 0, but with heavy tails that allow some terms to be regularised far from zero (Fig. S1). Furthermore, the regularised horseshoe prior has been successfully implemented to estimate interspecific interactions of two focal plant species with other species in their communities (Weis-Lehman et al. 2021). Here we apply the regularised horseshoe prior to estimate a full interaction matrix. See appendix S2 for further justification for our approach, and details about this prior and its implementation.

Because the main diagonal elements of the interaction matrix **B** represent effects of competition within species, which may come from a different distribution than the between-species effects, we estimate those terms using a conventional Gaussian prior (Fig. 2).

### Model fitting to data

To estimate the relative importance of interspecific interactions, intraspecific density dependence, and response diversity, we fitted our model (Fig. 2) to the fish count data. We also fitted a version with reef level random effects (see appendix S3 and Fig. S2).

We used the software program Stan (Stan Development Team 2023) which uses Hamiltonian Monte Carlo (HMC) sampling, because this approach provides a much greater range of tools to detect potential model pathologies that are not available for other MCMC algorithms, such as Gibbs samplers (Betancourt 2016; Monnahan *et al*. 2017).

From the 208 species in the LTMP, we generated data subsets with 20 and 40 species, prioritizing species with the smallest proportion of zero counts observed across reefs and time (Table S1). Collectively, these represent 52.49% and 64.93% of the total number of observed individuals in the data, respectively. All counts were analysed at species level except *Ctenochaetus* species, whose counts were grouped at the genus level and analysed it as a pseudo-species, *Ctenochaetus spp*, due to the resemblance between the two occurring species in the GBR, *C. binotatus* and *C. striatus*. For each dataset we ran 4 chains, each with 10000 iterations, 5000 iterations as warm up and 5000 as sampling. This left 20000 samples in the posterior distribution of each parameter. We used weakly informative priors and prior predictive checks (see Fig. 2 and appendix S4), to ensure that posterior estimates were informed by the data. Model convergence was monitored by examining posterior chains and distributions, running 4 chains with different randomly chosen initial values, checking that the potential scale reduction factor (R-hat) was close to 1 for all parameters, and checking that the effective sample sizes were large (Fig. S3). Model fit was assessed by posterior predictive checks (Fig. S3). Model predictive accuracy and model selection was evaluated by leave one out cross validation (LOO-CV) (Vehtari *et al*. 2017), using the R package *loo* (Vehtari *et al*. 2023) (Appendix S5).

### Simulation study

To assess the robustness of the parameter estimates and inferences produced in our analyses, we fit our models to simulated data with known parameter values. Specifically, we wished to verify that our approach could accurately estimate the off-diagonals of the interaction matrix **B** and the covariance matrix Σ, and thereby successfully distinguish between covariances in abundance produced by species interactions versus correlated responses to environmental fluctuations. In the baseline simulation, we set interspecific interactions and environmental covariances to zero to determine, whether model fits would erroneously identify non-zero interactions or covariances. In other simulations, we simulated non-zero environmental covariances and set the interspecific interactions terms to zero; we simulated communities where a subset of interspecific interactions was non-zero but there were no environmental covariances; and we simulated communities with both non-zero and some interspecific interactions were non-zero. See Appendix S6 for further details.

## Results

### Model fit to LTMP data

Model fits indicated very weak interspecific interactions, relative to conspecific density-dependence, but high response diversity. For interspecific around 97% of estimated posterior means had magnitudes between −0.01 and 0.01, more than an order of magnitude smaller than the mean intraspecific density-dependence (Fig. 3). In contrast, intraspecific density dependence was detected in all species with mean *b_ii_* < 1 and centred around 0.86 (i.e., mean strength of density-dependence 1-0.86=0.14: Fig. 3).

**Figure 3.**
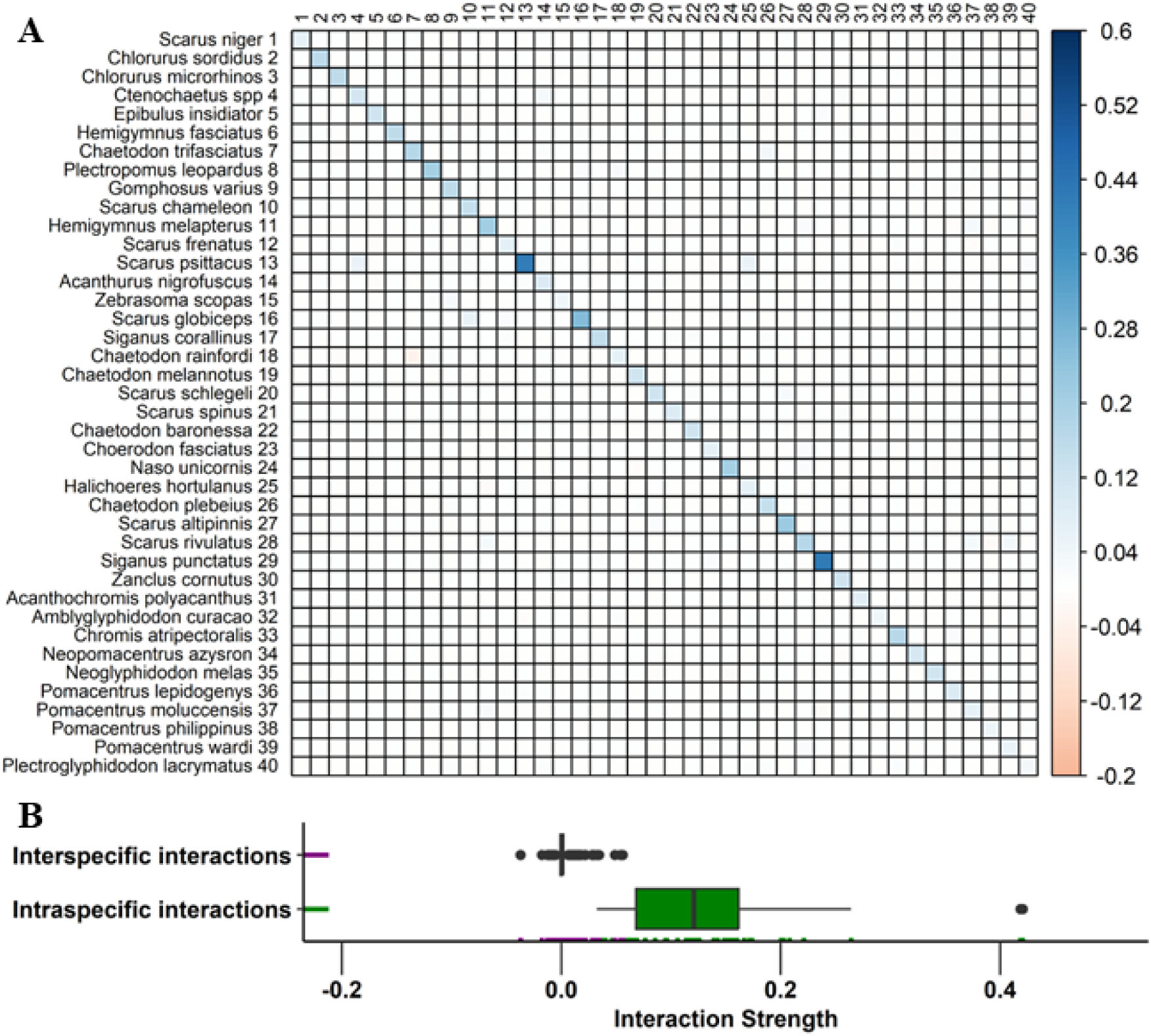
**(A)** Mean posterior estimates for the elements of the interaction matrix, B. **(B)** Distribution of posterior mean estimates for the elements of the interaction matrix. The diagonal elements are shown in the green boxplot (NB: the diagonal elements are presented here as 1-b_ii_ [i.e., 1 minus the diagonal element], such that zero implies density-independent growth, and positive values imply negative density-dependence). The off-diagonal elements are shown in the purple boxplot (NB: b_ij_ = 0 for no interaction; b_ij_ < 0 for negative effects (e.g. competition); b_ij_ > 0 for positive effects (e.g. facilitation)). The marks displayed along the vertical axis represent each of the estimated posterior means for the diagonal elements in green and off diagonal elements in purple.

In contrast to species interactions, species pairs exhibited a broad range of correlations in their responses to environmental fluctuations, with most weakly to moderately positively correlated, indicative of reasonably strong response diversity: most of the posterior mean estimates for the correlation values are between 0 and 0.5 (Fig. 4). The model with only 2 latent variables had the highest support. Model support decreased with increasing number of latent variables (Table S2) and the models with 10 and 12 latent variables did not converge. Model diagnostics indicated no convergence issues for the models with 2, 4, 6 and 8 latent variables. Thus, we selected the model with 2 latent variables for our analysis. Overall, however, the number of latent variables had a very small effect on the estimated interspecific correlations, with estimates remaining very similar as the number of latent variables increased from 2 to 12 (Fig. S4).

**Figure 4.**
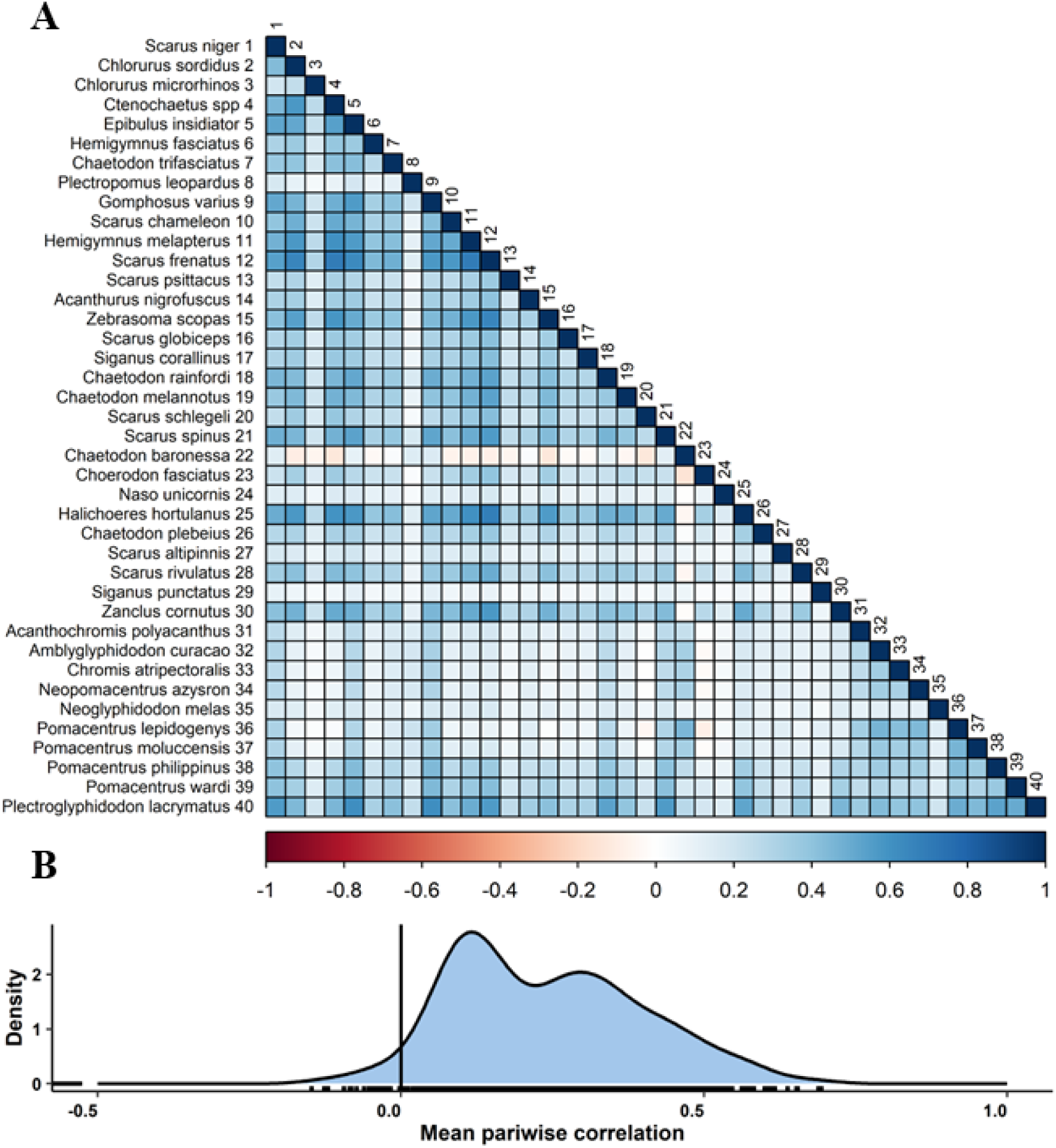
**(A)** Correlation plot showing the diagonal and the lower triangular elements of the correlation matrix, calculated from the variance-covariance matrix Σ as 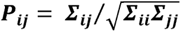. Each square shows the mean correlation estimate for a pair of species. **(B)** Density plot showing the distribution of posterior mean correlations. The black marks displayed along the horizontal axis represent each of the estimated posterior correlation means.

Overall, GBR fish assemblages exhibited similar magnitudes of variation in their species’ mean intrinsic growth rates (*CV**_a_*** = 0.58), intraspecific density dependence (*CV**_b_*** = 0.64) and sensitivity to environmental fluctuations (*CV*_σ_ = 0.59) (Fig. 5). The intrinsic growth rate parameter, *a*, varied among species but was consistently above zero (indicating capacity for recovery from low population density, i.e., persistence) with mean values from 0.03 to 0.5 (Fig. 5 panel A and B). Most species had mean intraspecific density dependence values (*1-b_ii_*) below 0.25 and above 0, with only 3 species having intraspecific density dependence values above 0.25 (Fig. 5 panel C and D), indicating weakly compensatory density-dependence (Thibaut & Connolly 2020). The standard deviations of the temporal variation in the density-independent growth rate (i.e., the square root of the diagonal of the variance-covariance matrix Σ) was right-skewed, with most species’ mean values close to the overall metacommunity mean value 0.54, and species-specific posterior means ranging from 0.19 to 1.45 (Fig. 5, E, F).

**Figure 5.**
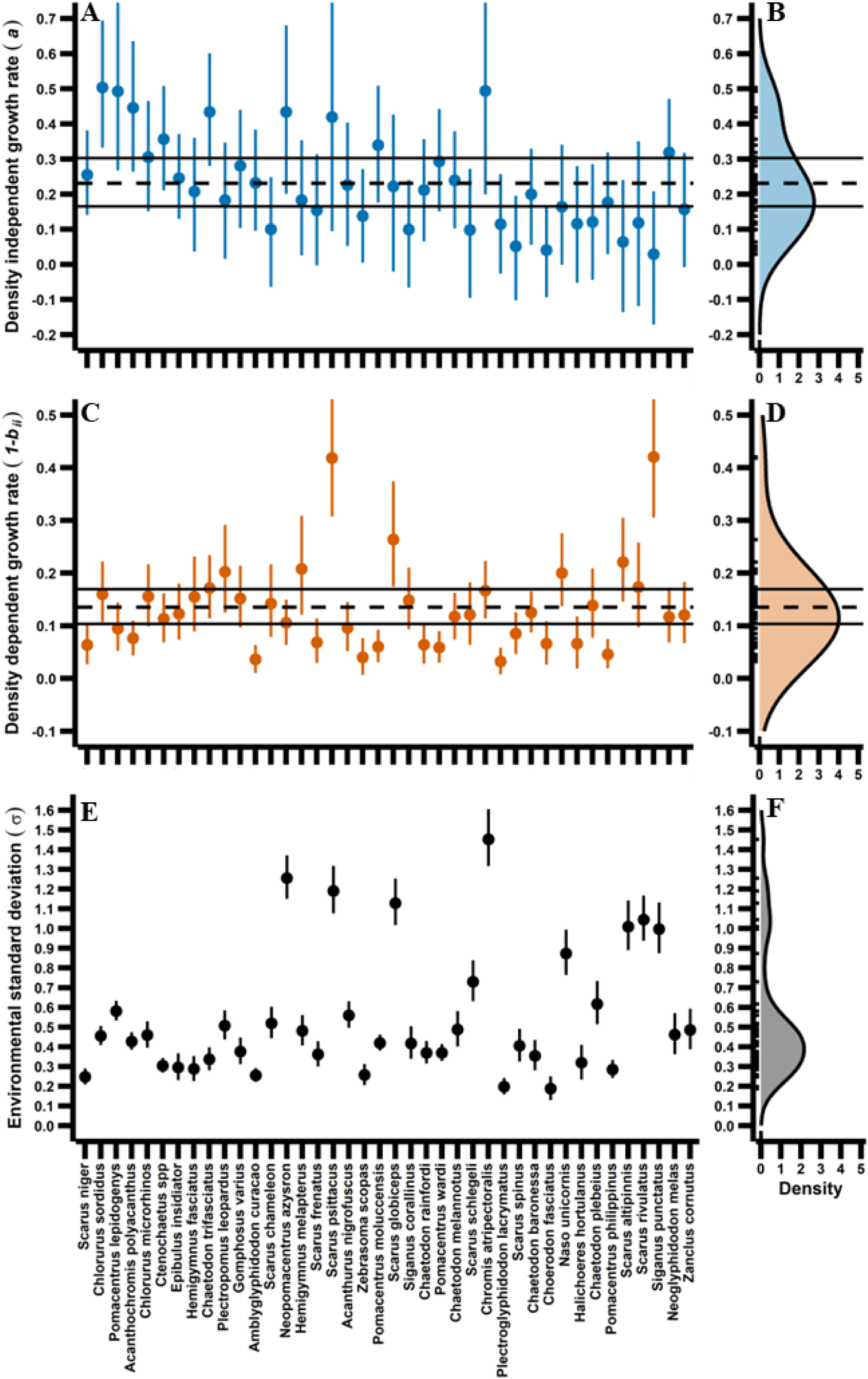
Interspecific heterogeneity in population-dynamic parameters. Panel **(A)** shows in blue each species posterior estimates for the density independent growth rate (i.e., intrinsic growth rate). Each dot represents a species posterior mean and the vertical lines represent the 95% credible intervals. Panel **(B)** shows the distribution across species of the posterior mean estimates from panel **(A)**. Each of the black lines displayed along the vertical axis in panel **(B)** correspond to each species posterior mean density independent growth rate (i.e., the dots in panel **(A))**. The dashed horizontal line in panels **(A)** and **(B)** show the higher hierarchical metacommunity mean (see Fig. 2 and eq. 1-3) and the two horizontal solid lines represent the upper and lower 95% credible intervals for that estimate. The same is displayed in orange for the species density dependent growth rate (i.e., intraspecific density dependence) estimates in panels **(C)** and **(D)**, and black for the species environmental standard deviation (i.e., standard deviation in temporal abundance) estimates in panels **(E)** and **(F)**. The species names on the horizontal axis for panels **(A)**, **(C)** and **(E)**, are sorted from the species with the lowest proportion of zero counts across reefs and years on the left, *Scarus niger*, to the species with the highest proportion of zero counts on the right, *Zanclus cornutus*.

We obtained consistent results regardless of whether models were fitted with or without reef-level random effects (Fig. S5). Surprisingly, we found evidence of very limited variability among reefs in intrinsic growth rates or intraspecific density dependence, for any species (Fig. S6, Appendix S3) with posterior distributions overlapping. Similarly, results are robust to the number of species included in the analysis. Fig. S7 compares parameter estimates from the model fits to 20 species and 40 species, for those species that appeared in both analyses: this shows that the model parameters were extremely similar in the two analyses (Fig. S7A, C, D). The interspecific species interaction parameters did appear to experience somewhat stronger shrinkage towards zero for the model with a higher number of species (FigS7B). However, most interactions were close to 0 in both analyses, and the few values farther away from zero did not show this shrinkage, indicating that the regularised horseshoe prior successfully allows nonzero interactions to escape the shrinkage to zero (also see “*Simulation study”* below).

### Simulation study

The simulation study indicated that the models successfully distinguished between zero and non-zero environmental correlations and interactions terms. Specifically, where these terms were zero, posterior means tended to be close to zero and had credible intervals encompassing zero. Conversely, when true parameter values were far from zero, posterior means and credible intervals correctly captured the direction and approximate magnitude of these effects, even when there were non-zero effects in off-diagonals of both the environmental correlation matrix and the interaction matrix (see Appendix S7 for detailed results).

## Discussion

We found that coral reef fish assemblages on the GBR exhibit a classically Gleasonian community structure (Gleason 1939), with highly heterogenous responses to environmental fluctuations and no evidence of interspecific interactions playing a strong role in the dynamics of species relative abundances. Conversely, these assemblages exhibited strong evidence of intraspecific density regulation, with detectable conspecific density-dependence for all species. There was, moreover, a high degree (i.e., CV ∼ 0.5 or larger) of demographic heterogeneity among species in density-independent and density-dependent components of population growth, as well as in their sensitivity to environmental fluctuations (i.e., the diagonal terms in the variance-covariance matrix Σ). This suggests that species differences in both deterministic (i.e., persistent through time), and stochastic (i.e., responses to environmental fluctuations) components structure patterns of commonness and rarity in fish communities, contrary to biodiversity theory based on approximate ecological equivalence, such as the lottery hypothesis (Sale 1977, 1978), and Neutral Theory of Biodiversity (Hubbell 2001). It also suggests more complex interspecific heterogeneity than assumed in at least some tractable alternative theories of biodiversity (Kalyuzhny *et al*. 2015; Engen & Lande 1996).

Our study also demonstrated successful fit of a multivariate community-dynamics model to time series data and accurate estimation of interspecific interactions and covariances in responses to environmental fluctuations even over a decadal time frame (∼10 years). We achieved this by leveraging the high degree of spatial replication, along with the dimension-reduction techniques of the regularised horseshoe prior and a latent environmental variable approach. In particular, using the regularized horseshoe prior, in lieu of a discrete mixture modelling approach (Mutshinda *et al*. 2009), allowed us to use Hamiltonian Monte Carlo and thereby exploit the more extensive model diagnostics available for such models, relative to alternatives such as Gibbs samplers, for which model pathologies can occur without tools to identify them (Betancourt 2016; Monnahan *et al*. 2017). These diagnostics, along with our simulation study, indicated that our approach yielded robust estimates of community dynamics parameters for species-rich communities, and can do so even for relatively short time series when sufficient replication is available.

Our finding that interspecific interactions were negligible seems at odds with classic and recent studies documenting strong interspecific interactions in nature (Connell 1961; Krebs *et al*. 2017; Paine 1966; Stenseth *et al*. 1997). In reef fish communities in particular, interspecific interactions have been considered to play a major role in structuring reef fish communities based on both field experiments (Jones 2005; Robertson 1996; Shulman 1985) and observational studies (Ebersole 1977). However, other reef fish studies have argued that interspecific interactions play a more limited role (Choat & Bellwood 1985; Mumby & Wabnitz 2002; Robertson & Seldon 1979). A feature of these past reef fish studies is that they have focused on small scales, where particular interactions are frequent, and on response variables whose changes can be measured readily at such scales, such as home range size or location (Jones 2005). However, such effects may be restricted in time or space, and thus have effects that do not scale up to the population level. For instance, strong heterospecific aggression does not necessarily translate into competitive release at the population level, when dominant competitors are removed (Blowes *et al*. 2017).

In contrast to interspecific interactions, we found strong evidence for compensatory intraspecific density dependence in fish assemblages, with mean values more than an order of magnitude larger than those estimated for interspecific interactions, and similar in magnitude as earlier estimates for insects, fishes, birds and mammals from analysis of single-population time-series (Thibaut & Connolly 2020) (Fig. S8). This finding of much stronger intraspecific than interspecific density dependence is consistent with a recent meta-analysis of pairwise interactions in plant studies (Adler *et al*. 2018). Similar findings also have been obtained from time series analysis of temperate vertebrate and invertebrate communities (Mutshinda *et al*. 2009; Ovaskainen *et al*. 2017b; Sandal *et al*. 2022). However, it is important to note that our study, like those cited above, focuses on interactions within a particular taxonomic group. Thus, the phenomenology of strong intraspecific against weak interspecific interactions could emerge from the action of species-specific natural enemies from other taxonomic groups, such as parasites and viruses, whose effects could become stronger as species become more abundant.

Our finding of large (CV > 0.5) and comparable degrees of heterogeneity in temporal average density-independent growth rate (*a_i_*), intraspecific density-dependence (*b_ii_*), and sensitivity to environmental fluctuations (σ*_ii_*), is inconsistent both with the demographic equivalence assumed by neutral theory of biodiversity (Hubbell 2001), and with the Engen model (Engen & Lande 1996; Engen *et al*. 2002), which assumes that interspecific heterogeneity is confined to the mean intrinsic growth rate term. The substantial variation in pairwise correlations in environmental responses is also inconsistent with this latter theory’s assumptions. However, the partitioning of variance in species abundances between deterministic and stochastic component, the principal application of this theory (Engen *et al*. 2002; Engen *et al*. 2011; Solbu *et al*. 2018; Bellier *et al*. 2022), appears to be fairly robust, at least for our system, as calculating these component using our full fitted model yields very similar variance proportions to a previous application of the Engen model (Tsai *et al*. 2022; Appendix S8, Fig. S9).

There has been considerable debate about the extent to which variation in the strength of conspecific negative density-dependence (CNDD) can explain variances in abundance in tropical forest assemblages. CNDD has been reported to be stronger for rare species than for common species (Comita *et al*. 2010; Johnson *et al*. 2012; LaManna *et al*. 2017; Mangan *et al*. 2010), independent of abundance (Chen *et al*. 2019; Fricke & Wright 2017), or stronger for common than rare species (Zhu *et al*. 2015). However, these approaches rely on surrogate measures of fitness, such as the relationship between seed abundance and seedling abundance, to estimate CNDD, and it has been suggested that such approaches can bias estimates of CNDD upwards, more so for common than rare species (Detto *et al*. 2019; Hülsmann *et al*. 2021; but also see LaManna *et al*. 2021). Our study is unaffected by this artefact, since it relies on time-series data, and we found that more abundant species indeed tend to experience less density-dependence than rarer species, a pattern that remains even when accounting for potential effects of body size (Fig. S10; see also Rovere & Fox *et al*. 2019; Yenni *et al*. 2017 for other examples using analysis of population time series). However, the relationship is somewhat weak, indicating that the variability in species intraspecific density dependence likely underlies a relatively small proportion of the abundance variation observed in this system.

In contrast to CNDD and species interactions, patterns of covariation in species’ responses to environmental fluctuations have received much less attention, either theoretically or in previous analyses of community time series (Mutshinda *et al*. 2009; Ovaskaninen *et al*. 2017b; Sandal *et al*. 2022). Our finding that a relatively small number of latent variables (as few as two, Fig. S4) captures the overall pattern of variances and covariances in response to environmental fluctuations suggests that a relatively small set of common drivers (or multiple drivers whose correlated dynamics produce relatively few important axes of variation) explains much of the environmentally-induced variation in population fluctuations in this system.

Additionally, the weak to moderate correlations in environmental responses indicate a relatively high degree of response diversity (i.e., asynchrony in population fluctuations) in this system, and thus a reasonably strong portfolio effect (Tilman *et al*. 1998; Elmqvist *et al*. 2003; Thibaut & Connolly 2013). Because interspecific interactions were negligible, this response diversity (i.e., heterogeneity in species responses to environmental fluctuations) is the overwhelming driver of community asynchrony for fishes on the GBR. In this respect, our findings are similar to an earlier analysis at the functional group level, which found similarly weak to moderate positive correlations between functional groups (Thibaut *et al*. 2012). Our results indicate that these conclusions also apply at the species level. However, a more detailed look at our results does reveal that environmentally-mediated correlations between species from the same trophic groups (Table S1) are slightly higher, on average, than those between species from different functional groups (Fig. S11A, B), and, similarly that more closely related species have slightly more positive environmental correlations than distantly related species (Fig.S11C, D, Fig. S12 and Appendix S9). However, these differences are small and explain very little of the overall variation in the structure of the environmental correlation matrix, suggesting that the factors driving these correlations are highly idiosyncratic and not strongly conserved phylogenetically, nor dependent on the nature of a species’ trophic role.

Species interactions have been hypothesized to play strong roles in the ecology and evolution of communities, particularly in the tropics, shaping phenomena from equatorward range limits (Darwin 1964; Paquette & Hargreaves 2021), to the latitudinal diversity gradient (Dobzhansky 1950; Schemske *et al*. 2009; Zvereva & Kozlov 2021), to macroevolutionary trends in taxonomic diversity, and ecosystem function (Vermeij 2019; Bush & Payne 2021). However, the curse of dimensionality has complicated assessing the role of such interactions at the whole-assemblage level in species-rich ecological communities. Our approach offers a way to confront community-dynamics models with time series from such high-dimensional systems, to rigorously explore the robustness of the models’ performance, and to infer the relative importance of such interactions, alongside other factors such as response diversity and other sources of demographic heterogeneity among species. For coral reef fishes, population regulation is driven overwhelmingly by intraspecific density-dependence, whereas interspecific interactions have negligible effects on population level dynamics, suggesting a very high degree of niche differentiation in this assemblage. We hope our work prompts similar analyses in other systems, to more comprehensively assess the factors that drive the dynamics of species abundances in high-diversity systems like coral reefs.

## Acknowledgments

We thank past and present members of the LTMP of Australian Institute of Marine Science, and the crews of the research vessels Harry Messel, Sirus, Lady Basten, Cape Fergunson and Solander. We also acknowledge the Great Barrier Reef Traditional Owners of the land and sea Country in which we work and pay our respects to their elders, past present and emerging. We thank HPC facilities of James Cook University for computation resources. This research was supported by an AIMS@JCU scholarship to AMR, by the Smithsonian Tropical Research Institute and by the Australian Research Council Centre of Excellence Program (grant number CE140100020).

## Author contributions

ARM and SRC conceived this study. MJE supervised and provided the LTMP data. ARM performed the modelling work and analysed the model output with help of SRC. ARM and SRC wrote the first draft of the manuscript and MJE contributed substantially to revisions.

