## Supplementary materials for "High response diversity and conspecific density-dependence, not species interactions, drive dynamics of coral reef fish communities"

**Appendix S1: Latent factor approach and identifiability**

A factor analysis approach applied to our multispecies model (eq. 1 and 2) can be formulated as

${log(\boldsymbol{\mu}}_{\boldsymbol{t}})\sim MVNormal(\boldsymbol{a+B log(}\boldsymbol{\mu}_{\boldsymbol{t-1}}\boldsymbol{)}+\boldsymbol{\Lambda} \boldsymbol{\theta}_{\boldsymbol{t}}, \boldsymbol{\Psi})$ (S1)

$\boldsymbol{\theta}_{\boldsymbol{t}}\sim MVNormal(0, \boldsymbol{\Phi})$	(S2)

where *log***(μ_t_)** is a *S*-dimensional vector of observable log-abundances that follows a multivariate normal distribution with a vector of means (***a* + B log(μ_t-1_) + Λθ_t_)**, and variance-covariance matrix **Ψ*.* θ_t_** is a D-dimensional vector of latent variables for time *t*, *t =* *1,…,n*. Thus, the matrix **ϴ** has *D x T* dimensions and **Φ** (*D x D*) is the covariance matrix of the latent variables. **Λ** is an *S x D* matrix of factor loadings. Here the latent variable *θ_kt_* (element from matrix **ϴ**) can be interpreted as the value of an unmeasured environmental covariate *k* at time *t* and *λ_jk_* (element from matrix **Λ**) is the response of species *j* to the unmeasured variable *k*.

To ensure identifiability of the variance parameters, the residual errors are assumed to be independent, thus **Ψ** is a diagonal matrix (Howe 1955; Lord and Novick 1968; Harvey 1990; Peeters 2012; Erosheva & Curtis 2017). Given that, the total covariance matrix becomes **Σ = ΛΦΛ’+Ψ**. Furthermore, the latent variables can be marginalized out and the model becomes

${log(\boldsymbol{\mu}}_{\boldsymbol{t}})\sim MVNormal(\boldsymbol{a+B log(}\boldsymbol{\mu}_{\boldsymbol{t-1}}\boldsymbol{)}, \boldsymbol{\Lambda\Phi\Lambda'}+\boldsymbol{\Psi})$ (S3)

However, a computational problem with this approach is that the factor analysis model is inherently unidentified. Both the latent variables and the factor loadings are unknown, and they have to be estimated. Given **Ψ**, two different sets of **Λ** and **Φ** can generate the same covariance structure **Σ** (Peeters 2012). Therefore, **Λ** and **Φ** need to be constrained to ensure that all chains converge to the same modes. There are multiple sets of constraints in the literature that ensure statistical identifiability of the model (Farouni 2015; Peeters 2012; Merkle and Wang 2018). In our case, the set of constraints that ensured model convergence were:

- **Φ** is a positive definite matrix such that Diag(**Φ**)=**I** (i.e., is the identity matrix).
- Fix the upper triangular elements of the **Λ** matrix to 0.
- Constrain the diagonal elements of the **Λ** matrix to be positive.
- The resulting **Λ** matrix is full rank.

Even with these constraints, identifiability issues still can arise. Therefore, we used a parameter expansion approach (Gelman 2004; Ghosh and Dunson 2009; Merkle and Wang 2018) to address identification issues not solved by the constraints above. Parameter expansion involves a *working model* with the unidentified parameters:

${log(\boldsymbol{\mu}}_{\boldsymbol{t}})\sim MVNormal(\boldsymbol{a+B log(}\boldsymbol{\mu}_{\boldsymbol{t-1}}\boldsymbol{)}, \boldsymbol{\Lambda}^{*}\boldsymbol{\Phi}\boldsymbol{\Lambda}^{*}'+\boldsymbol{\Psi})$ (S4)

where **Λ**^*^ is a *S x D* working factor loadings matrix having the constraints set above. This model is fitted to the data. At each iteration of the MCMC algorithm, we transformed the sampled parameters to a corresponding *inferential model* with uniquely identified parameters by using the following transformation:

$\lambda_{j,k}\boldsymbol{=}\left\{ \begin{aligned} {\lambda_{j,k}}^{*}{\varphi_{k,k}}^{1/2} if {\lambda_{k,k}}^{*}>0 \\ {{-\lambda}_{j,k}}^{*}{\varphi_{k,k}}^{1/2} if {\lambda_{k,k}}^{*}<0 \end{aligned} \right.$	(S5)

where *λ_j,k_^*^* is the *j^th^* and *k^th^* element from matrix **Λ**^*^ and *φ_k,k_* is the *k^th^* diagonal element of the matrix **Ψ**. This transformation combined with the set of constrains above made the chains of sampled parameters converge.

**Appendix S2: The regularised horseshoe prior**

Species interaction matrices have been estimated using a latent variable approach akin to that described in Appendix S1 for the variance-covariance matrix (e.g., Ovaskainen *et al.* 2017; Sandal *et al.* 2022). However, while in the context of the variance-covariance matrix the latent variables can be readily conceptualized as unmeasured environmental drivers whose fluctuations drive species’ responses, this has a less ready interpretation in the context of the interaction matrix (at least to us). In contrast, a discrete mixture approach (where a given interaction term has some probability of being drawn from a distribution with all of its probability mass at zero, and with one minus that probability is drawn from a continuous distribution that allows nonzero values; Mutshinda *et al.* 2009; but also see *sparse interactions* model in Ovaskainen *et al.* 2017 and in Sandal *et al.* 2022), is consistent with our conceptualization that, in a high-diversity community, most species pairs have negligible interactions but some are non-negligible. However, implementing a discrete mixture would have required use of a Gibbs sampler for MCMC fitting, which do not provide the extensive model diagnostics to identify potential pathologies (e.g., divergent transitions), as newer approaches such as Hamiltonian Monte Carlo (HMC) (Monnahan *et al.* 2017), even though such pathologies affect both HMC and Gibbs samplers (Betancourt 2016). Therefore, to be able to employ HMC, we instead used the regularized horseshoe prior, a continuous distribution that can also be employed in contexts where many parameters are very close to zero, but some may be far from zero (Piironen & Vehtari 2017).

To understand how the regularised horseshoe prior works, consider a Gaussian linear regression model

$y_{i}=\boldsymbol{\beta}\boldsymbol{x}_{\boldsymbol{i}}+ \varepsilon_{i}, \varepsilon_{i}\sim N\left( 0, \sigma^{2} \right), i=1,\ldots,n,$	(S6)

where **x_i_** is a vector of covariates for *i*, **β** is a vector of regression coefficients and *σ* is the residual error. The horseshoe prior (Carvalho *et al.* 2009) for **β** is:

$\beta_{j} \sim N(0, \tau^{2}{\lambda_{j}}^{2})$	(S7)

$\lambda_{j}\sim{Cauchy}_{(0,\infty)}\left( 0, 1 \right), j=1,\ldots,D$ (S8)

where *β_j_* is a weight parameter in the model. *τ* is the global hyperparameter that shrinks all the parameters towards zero, while the heavy-tailed half-Cauchy prior for the local hyperparameters *λ_j_* allows some *β_j_* to escape the shrinkage. The horseshoe prior has two shortcomings (Piironen and Vehtari 2017). First, if the parameters far from zero are weakly identified they can generate an unstable MCMC sampler**.** Second, there has been a lack of consensus on how to carry out inference for the global parameter *τ*. Piironen and Vehtari (2017) formulated the regularised horseshoe prior that addresses these two issues**:**

$\beta_{j} \sim N\left( 0, \tau^{2}\tilde{\lambda}_{j}^{2} \right),$	(S9)

$\tilde{\lambda}_{j}^{2}=\frac{c^{2}\lambda_{j}^{2}}{c^{2}{+\tau^{2}\lambda}_{j}^{2}} ,$	(S10)

$\lambda_{j}\sim C^{+}\left( 0, 1 \right), j=1,\ldots,D$	(S11)

where $C^{+}\left( 0, 1 \right)$ denotes a half Cauchy distribution truncated at 0 and *c*>0. If the true regression coefficient is close to zero, the local parameter *λ^2^_j_* will not be large enough to allow the estimated parameter *β_j_* escape the shrinkage towards zero by the global parameter τ*^2^*. Therefore, the term *τ^2^λ^2^_j_* will approach zero in the denominator in eq. S10, and therefore, $\tilde{\lambda}_{j}^{2}$will approach $\lambda_{j}^{2}$. In that case, *β_j_* will follow a normal distribution with mean zero and variance close to zero (eq. S9). In contrast, if *β_j_* is non-zero, the local parameter *λ^2^_j_* will allow the parameter *β_j_* escape the shrinkage towards zero by the global parameter τ^2^. Therefore, τ^2^λ^2^_j_ will be non-zero in the denominator of eq. S10 and if τ^2^λ^2^_j_ is much larger than *c^2^*, in the denominator of eq. S10 $\tilde{\lambda}_{j}^{2}$will approach ${c^{2}}/{\tau^{2}}$. Thus, *β_j_* will follow a normal distribution with mean zero and variance *c^2^*. In essence, when the data suggests values close to zero, the regularised horseshoe prior will behave like the original horseshoe prior and shrink the parameter towards zero. On the other hand, when the data supports for values far from zero, the regularised horseshoe prior will still regularize these large coefficients according to a Gaussian distribution with mean zero and variance *c^2^*, while the original horseshoe prior does not regularise large signals.

The global hyperparameter *τ* is defined as

$\tau\sim N^{+}\left( 0, \tau_{0}^{2} \right)$	(S12)

$\tau_{o}= \frac{p_{0}}{P-p_{0}} \frac{\sigma}{\sqrt{n}}$	(S13)

where $N^{+}\left( 0, \tau_{0}^{2} \right)$ denotes a half-normal distribution truncated at zero and with variance *p_0_* is a prior guess for the number non-zero parameters, *P* is the total number of regression parameters to be estimated, *σ* is termed the “noise level”, and *n* is the sample size.

In this study, we extend application of the regularized horseshoe prior from the univariate case (a single vector of observations) to the case of a multivariate response (i.e., a matrix of abundances of multiple species across multiple years). Specifically, we model the off diagonals of the interaction matrix **B**, b*_ij_*, in a fashion analogous to eq. S9-S13:

$b_{i,j} \sim N(0,\tilde{\lambda}_{i,j}^{2}\tau^{2})$	where *i ≠ j* (S14)

$\tau\sim N^{+}(0, \tau_{0}^{2})$	(S15)

${\tau_{0}}= \frac{p_{0}}{P-p_{0}} \frac{\sigma_{hyper}}{\sqrt{n}}$	(S16)

${\tilde{\lambda}_{i,j}^{2}}= \frac{c^{2}\lambda_{i,j}^{2}}{c^{2}+ \tau^{2}\lambda_{i,j}^{2}}$	(S17)

$\lambda_{i,j}= {local1}_{i,j}\times\sqrt{{local2}_{i,j}}$	(S18)

${local1}_{i,j}\sim N^{+}(0, 1)$	(S19)

${local2}_{i,j}\sim\mathrm{InvGamma}(1.5, 1.5)$	(S20)

We set the hyper-parameter *P* as the number of off-diagonal elements in **B**, and *n* as the total number of observations (i.e., abundance of each species, including 0 abundance, at each year at each reef). In the standard regression context (eq. S6), the “noise level” hyper-parameter *σ* (eq. S13) is related to the residual variance. In our case, however, the process error depends on the environmental covariance matrix (and thus can vary among species), and the observation error is assumed to be Poisson. We chose to calculate the noise level hyper-parameter in eq. S16 as $\sigma_{hyper}= mean(\sqrt{diagonal(\boldsymbol{\Sigma})})$. This choice is somewhat arbitrary, and thus may not be optimal, but simulations show that the between species interactions parameters are retrieved correctly.

**Appendix S3: Random effects**

We used a multilevel structure in the model to account for the variability in the intrinsic growth rate and the within-species density dependence at the species level and at the reef level (Fig. S2 blue box). Therefore, *a_i,r_* is the intrinsic growth rate of species *i* at reef *r*, and it follows a normal distribution with mean $\bar{a}_{i}$ and standard deviation $\sigma_{ai}$ for species *i* (see Reef level box in Fig. S2). Furthermore, all species mean intrinsic growth rates, $\bar{a}_{i}$, are drawn from a normal distribution with mean $\bar{a}$ and standard deviation of $\sigma_{a}$(Fig. S2). The mean $\bar{a}$ has a normal prior distribution with mean 0 and standard deviation of 1 (Fig. S2). This multilevel structure sets 3 levels: metacommunity level, species level and reef level. The metacommunity level estimates the mean intrinsic growth rate across species, for the metacommunity as a whole. The species level characterizes the among-species variation in intrinsic growth rates, and the reef level allows for spatial variability in the intrinsic growth rate of each species across reefs. The same multilevel structure was implemented for the within species density dependence (see Fig. S2). We only modelled the diagonal of the interaction matrix with this multilevel structure, due to the high computation cost and lower parameter identifiability of estimating a full interaction matrix for each reef replicate. Fig. S2 shows the prior choices for different parameters in the multilevel part of the model.

Our initial attempt to fit this model yielded low effective sample sizes, particularly when including Pomacentrid species in the model. To ensure model convergence we implemented three changes. Firstly, we used somewhat tighter priors on the standard deviation terms ($\sigma_{ai}$, $\sigma_{bii}$, $\sigma_{a}$, and $\sigma_{b}$, see Fig. S2; these priors were still vague enough to not bias the posterior inference). Secondly, we ran the model for 20000 iterations, 10000 iterations as warm up and 10000 as sampling, yielding 40000 samples across the four chains in the posterior distribution of each parameter. Finally, we fitted the model to the 20 most common non pomacentrid species out of the 40 in our analysis in the main text.

**Appendix S4: Prior choice and prior predictive check**

For the intrinsic growth rate and the intraspecific density dependence, we chose vague priors to ensure that the likelihood was driving the posterior estimation (see Fig. 2). Specifically, for the intraspecific density dependence, the prior choice was vague enough that covered values above zero, which would indicate negative density-dependence, and values below zero, indicating positive density-dependence. Our posterior estimates were between 0 and 1 indicating compensatory negative density-dependence, and the posterior distribution was much narrower than the prior (compare “prior” and “posterior” at the top of Fig. S8).

For the factor loadings, we implemented wide priors according to the literature (Erosheva & Curtis 2017; Farouni 2015; Ghosh & Dunson 2009) (See methods and appendix S1 for constraints implemented in the analysis).

For the regularised horseshoe prior, we implemented similar hyperprior choices that Piironen and Vethari implemented in their study (2017). We set 𝜈_local_ and 𝜈_global_ to 3 and 1 as these values produced fewer convergence problems when the model was fitted to the LTMP data.

We also implemented prior predictive checks, which in essence involve simulating data from the priors and investigating the consistency of the model predictions with domain knowledge, given our prior choice. Given the complexity of our model, we partition the prior predictive check into different parts: Firstly, we simulated 1000 communities of 20 species each for 1000 years with the priors from Fig. S2 for the intrinsic growth rate and the intraspecific density dependence. We assumed no interspecific species interactions, no covariance in species environmental fluctuations, and we set the sensitivity to environmental fluctuations (the diagonal elements of the variance-covariance matrix) to 0.1 for all species. The simulated abundances went from zero abundance to the maximum real value a 64-bit machine can handle (1.797693e+308), indicating that the priors were vague enough to produce unrealistically high and low values (Fig. S13). Secondly, we simulated 1000 variance-covariance matrices for 20 species and 2 latent variables to estimate the distributions of correlations and standard deviations in abundances that the latent variables produced given our prior choice (Fig. 2). The correlation values had a symmetrical u-shaped distribution centred around 0 with values ranging between -1 and 1, and the temporal standard deviations values were centred around 12.32 with values ranging close to 0 to close to 60 (Fig. S14). Both distributions were far broader that, and extended both above and below, the corresponding posteriors (see Results), indicating that they are unlikely to have materially informed the posteriors.

Finally, we implemented 1000 simulations to estimate the distribution of interspecific interactions terms that the regularised horseshoe prior produced (Fig. S15). We assumed that *n* in eq. S16 is 10000, *P* is assumed to be calculated for 20 species (i.e., off-diagonal elements of 20-by-20 interaction matrix), and thus is 380, *p0* was set to 10 and the σ_hyper_ value was close to the mean σ_hyper_ value from the model fit to the LTMP data (i.e., square root of the mean of the diagonal of **Σ** in eq. 5). Given our prior choice, the simulated parameters were centred around 0 with a range between -15 to 15, approximately, again orders of magnitude broader than the distribution of the posterior estimates at estimates.

**Appendix S5: Cross-validation and model selection**

To use the *loo* package, the model written in Stan needs to calculate and store the pointwise log-likelihood using the posterior. Then, the function “loo” computes the efficient PSIS-LOO approximation to exact LOO-CV (Vehtari *et al.* 2017, 2022; also see vignette by Vehtari & Gabry 2023). In our case, we have at least one parameter for each observation (i.e., the model estimates the expected abundance for species *i* at time *t*, *μ_i,t_*, for each observed count, *y_i,t_*, in the data eq. 1-4). Therefore, when one observation is removed, the posterior for the corresponding parameter changes significantly and the PSIS cannot approximate well the exact LOO (Vehtari 2017). For univariate cases, adaptative quadrature can improve stability (Vehtari *et al.* 2016; also see *Poisson model with “random effects” and integrated LOO* in Vehtari 2017). In essence, this approach consists in integrating out the problematic parameter to estimate the pointwise log-likelihood. However, Stan can only integrate over one dimension and nested integration would be very slow when compared to MCMC methods for more than two parameters. Instead, we calculated the likelihood of each species by using the function *dpoilog* from the *poilog* package (Grøtan & Engen 2022). Then, we calculated the log of the sum of all species likelihood at each time step to obtain the pointwise log-likelihood, which we used with the package loo to estimate LOO-CV and perform model selection (See Table S2). Note that this approach ignores the species environmental correlations as *dpoilog* is providing the likelihood for the univariate Poisson lognormal distribution.

**Appendix S6: Simulation scenarios**

Our objective is not to broadly examine the statistical performance of the model for a vast combinations of parameter values, but rather to verify that the latent variables and the regularised horseshoe prior can retrieve the off diagonals of the interaction matrix **B** and the covariance matrix **Σ** with minimal bias for data sets with the structure of the LTMP fish data. Therefore, we used similar parameter values to the estimated posterior distributions obtained from model fitting to the LTMP data (See results). To do so we simulated different scenarios for 20 species on 41 reefs for 1000 years. To use the same time series length, we only analysed the last 11 years in the simulations. Also, if a model fit did not converge we simulated a new community with the same parameters. Here we present simulations for which the model does converge.

- *Baseline*: In this simulation we wanted to test whether the model recovers the interspecific interactions and species correlations in their responses to environmental fluctuations when they are zero in generating process simulating the data. The rest of the parameters were similar to the parameter estimates from the model fit to the LTMP data. The species intrinsic growth rate was drawn form a truncated normal distribution (from 0 to ∞) with mean 0.2 and standard deviation 0.15. The species intraspecific density dependence was drawn form a truncated normal distribution (from -∞ to 1) with mean 0.85 and standard deviation 0.05, the species standard deviation to environmental fluctuations was drawn from a truncated normal distribution (from 0 to ∞) with mean 0.5 and standard deviation 0.2. If the regularised horseshoe prior is correctly implemented, it should shrink all the off-diagonal elements of the interaction matrix B to zero, with their mean values close to zero and their credible intervals containing zero. Similarly, the latent variable approach should give correlation values with mean values close to 0 and credible intervals containing zero.
- *Between species interactions*: Differently to the *baseline* simulation, here we wanted to test whether the model recovers the interspecific interactions when present and while correctly estimating the uncorrelated responses in species responses to environmental fluctuations. Specifically, that the regularised horseshoe prior shrink towards zero the pairwise interactions that are zero while still correctly identifying and estimating the parameters that are nonzero. We created an interaction matrix **B** where we arbitrarily chose about 10% of the interspecific interaction parameters to be non-zero, while the rest of the off diagonals were set to 0. Also, we estimated the dominant eigenvalue of the interaction matrix to ensure the stability of the population as well as ensuring that no species had exponential temporal trajectories. The diagonal elements were drawn from a normal distribution with mean 0.85 and standard deviation 0.001. This was done to ensure that the simulations were stable (i.e., the community fluctuated around a long-term average state). The rest of the model parameters values were the same as the *Baseline* scenario. We also fitted the model with *p0* = 10, *p0* = 50 and *p0* = 100 (prior guess for the number of non-zero parameters) to investigate the sensitivity of the posterior parameter estimates for the interspecific interactions to *p0*.
- *Correlated responses to environmental fluctuations*: Here we wanted to test the converse of the previous scenario: that is, can the model correctly retrieve the nonzero correlations in species responses to environmental fluctuations while there are no interspecific interactions? The model should correctly assign all covariation in species abundances to nonzero elements in the species variance-covariance matrix **Σ**, while shrinking all the off-diagonals of the interaction matrix **B** towards zero. To do so, we generated a random correlation matrix by using the *randcorr* package (Makalic & Schmidt 2022). This produces a correlation matrix with most of the correlation values between -0.5 and 0.5, with its mean at 0. The rest of the model parameters values were the same as the *Baseline* scenario. Also, we fitted the model with different number of latent dimensions (*D*): *D* = 2, *D* = 4, *D* = 6, *D* = 8, *D* = 10, and *D* = 12.
- *Both*: In the last scenario we want to test if the model can distinguish between interspecific interaction mediated and environmentally mediated fluctuations in abundance. We implemented the interaction matrix and the correlation matrix from the *Between species interactions* and the *Correlations scenario*, respectively. The rest of parameters values were the same as the *Baseline scenario*. For simplicity, we fitted the model with *p0* = 10 and *D* = 2.

**Appendix S7: Simulation results**

For the baseline simulation in which there were no interspecific interactions or covariances in responses to environmental fluctuations, the model correctly identified this independence among species. Model posterior mean estimates for pairwise environmental response correlations tended to be very close to zero, and those species pairs for which posterior means were less close to zero had high uncertainty, such that all credible intervals encompassed zero (Fig. S16 Panel A). Similarly, posterior mean estimates for the interspecific interactions were close to and had credibility intervals that encompassed zero (Fig. S16 Panel B), generally with more precision than the estimated environmental response correlations.

For the simulation in which interspecific species interactions were present for a subset of species, but environmental correlations were absent, the model correctly identified the nonzero interaction terms without artefactually introducing additional nonzero interaction terms or environmental correlations. As with the baseline scenario, environmental correlation terms were small and had credible intervals encompassing zero (Fig. S17 Panel A). Similarly, posterior estimates of interactions terms for non-interacting pairs were all very close to zero, whereas estimates for the nonzero interactions were close to the true values, with some evidence of regularization (i.e., large positive interaction term estimates tended to be slightly less positive than the true values, and large negative interactions were estimated to be slightly less negative than the true values: Fig. S17 Panel B). Also, the interspecific interaction parameter estimates were identical regardless of the prior guess of non-zero parameters, *p0* (Fig. S18).

In the simulation scenario in which species variability in abundance was due to environmental correlations only (i.e., all interspecific species interactions were zero), the model correctly identified the negative, positive and near-zero environmental correlation values without artefactually estimating any of species’ interactions as different from zero. Posterior estimates for the environmental correlations were close to the true values (Fig. S19 Panel A). As with the baseline scenario, interspecific interactions terms were close to and had credibility intervals that encompassed zero (Fig. S19Panel B). Also, the higher the number of latent variables, the closer to the true parameters implemented in the simulation (Fig. S20).

For the simulation including both interspecific species interactions and environmental correlations, the model was able to discern between interspecific interaction mediated and environmentally mediated fluctuations. The posterior estimates for the environmental correlations were close to the true values (Fig. S21 panel A) and the results were similar to the simulation scenario with only species correlations (compare with Fig. S19 Panel A). Although there were three non-zero interactions that were shrunk toward 0 by the model, the regularised horseshoe prior captured well the rest of interaction parameters implemented in the simulation, including the non-zero terms (Fig. S21 panel B).

**Appendix S8: Variance Partition of Relative Species Abundance**

Engen *et al.* (2002) derived an approach to explain the drivers of species abundances by partitioning the variance of relative species abundance. This theory was derived from a stochastic model that defines the dynamics of the log abundances by a continuous Ornstein-Uhlenbeck process

${dX}_{i}= (r_{i} - \delta X_{i})dt + \sigma_{e}dW_{i}$ (eq. S21)

$r_{i} \sim N(\mu_{r},\sigma_{r}^{2})$	(eq. S22)

where *X_i_* is the log abundance of species *i*. *r_i_* is the intrinsic population growth rate of species *i* (analogous the intrinsic growth rate *a_i_* in eq. 2) and it follows a normal distribution with mean *μ_r_* and standard deviation *σ_r_*. *δ* measures the strength of density dependence (analogous to intraspecific density dependence b_ii_ in eq. 1), but in this case it is constant across species. *σ_e_* scales the magnitude environmental fluctuations in the growth rate (and is also assumed to be the same for all species), and *dW_i_* models the fluctuations as standard Brownian motion, and thus there is no covariation in species responses to environmental fluctuations.

Given the model in eq. S21 and eq. S22 and its restrictive assumptions, the total variance (σ^2^_total_) in the log abundance among species can be decomposed into three additive components:

$\sigma_{total}^{2}= \frac{\sigma_{r}^{2}}{\delta^{2}} + \frac{\sigma_{e}^{2}}{2\delta} + \theta^{2} = V_{r}+V_{e}+V_{d}$ (eq. S23)

where ${{V_{r}=\sigma}_{r}^{2}}/{\delta^{2}}$ represents the variance in species log-abundances due to the variability in their intrinsic growth rates, ${{V_{e}=\sigma}_{e}^{2}}/{2\delta}$ represents the variance in species log abundance due to species’ response to environmental fluctuations, and $V_{d}=\theta^{2}$ represents the variance in species log-abundance due to residual variation caused by other processes such as overdispersion. These variance components can be estimated from the correlation coefficient (ρ_t_) of log-species abundance between two communities with a time lag of t as

$\rho_{t}= (\rho_{0}-\rho_{\infty})e^{-\delta t} + \rho_{\infty}$ (eq. S24)

where ρ_t_ can be interpreted as a measure of community similarity between species log-abundances at two different times. This is modelled as an exponential decay of the time lag t. δ is the strength of within species density dependence as in eq. S21. ${{\rho_{\infty}=(\sigma}_{r}^{2}}/{\delta^{2})/}\sigma_{total}^{2}$ = $V_{r}/\sigma_{total}^{2}$ represents asymptotic similarity. If all species in a community have the same growth rate as well as density-dependence (i.e., $\sigma_{r}^{2}=0$), this value will tend to zero. In contrast, for a community where environmental fluctuations play no role, and differences in species’ demographic parameters explain most of the variation in abundances, ρ_∞_ would be large. ${{\rho_{0}=(\sigma}_{r}^{2}}/{\delta^{2} +{\sigma_{e}^{2}}/{2\delta} )/}\sigma_{total}^{2}$ = ${(V_{r} + V_{e})/\sigma}_{total}^{2}$ is the intercept and it represents the theoretical expected correlation in species’ log abundances when the time lag is zero. This parameter would tend to unity in the absence of demographic stochasticity, sampling error, and overdispersion.

With estimates for ρ_∞_, ρ_0_ and δ, the three additive components of the total variance can be calculated as

$V_{r} = \rho_{\infty}\sigma_{total}^{2}$	(eq. S25)

$V_{e} = (\rho_{0}-\rho_{\infty})\sigma_{total}^{2}$ (eq. S26)

$V_{d} = {1 - \rho}_{0}\sigma_{total}^{2}$	(eq. S27)

Analytically, eq. S21 and eq. S22 show that each species’ abundance fluctuates around their carrying capacity, $e^{\frac{r_{i}}{\delta}}$ and both the carrying capacities and the abundances themselves follow a lognormal distribution among species. Therefore, this model could be considered analogous to the discrete Gompertz model (Ives *et al.* 2003), which has the same analytical result for species carrying capacities.

This method of variance partitioning is derived from the model in eq. S21 and eq. S22, and thus entails the assumptions of that model. In contrast, our model from eq. 1-3 (see Fig. 2) is much more flexible with respect to species interactions: intraspecific density-dependence can vary among species, and interspecific interactions can be present. In addition, our model allows species differences in the magnitude of environmentally induced fluctuations, and nonzero covariances in species’ response to environmental fluctuations. Therefore, to estimate the proportional variance under these less restrictive assumptions we used the general analytical solutions (Ives 2003) of the multivariate Gompertz model to estimate the environmental variance and variance in equilibrium abundance, following Tsai et. al (2022):

$\nu_{r} = var(log \hat{N}) = var[{(I-B)}^{-1} A]$	(eq. S28)

$\nu_{e} = \bar{V}_{e} =avg[{(I-B⦻B)}^{-1} vec(\Sigma)]$	(eq. S29)

where 𝜈_r_ is the variance among species in equilibrium log-population sizes, and 𝜈_e_ represents the average species-level variance of log-abundance due to environmental fluctuations. In eq. S28, $\log\hat{N}$ represents the species’ abundances at equilibrium states, “var” represents the variance operator, B is the interaction matrix (as in eq. 2), and A is a vector of intrinsic growth rates (as in eq. 3). In eq. S29, $\bar{V}_{e}$ is the average of the diagonal of the environmental variance-covariance matrix at stationary states, “avg” represents the arithmetic mean function, and “diag” and “vec” are the diagonal and vectorization operators. The symbol $⦻$ denotes the Kronecker or direct matrix product. Therefore, the proportional variance due to niche differences and different responses to environmental fluctuations can be defined as:

$P_{niche} = \frac{\nu_{r}}{\nu_{r}+\nu_{e}}$ (eq. S30)

$P_{env} = \frac{\nu_{e}}{\nu_{r}+\nu_{e}}$	(eq. S31)

We found that our variance partition estimates from eq S30-S31 were similar to the estimates calculated using the Engen model (Engen *et al.* 2002) in reef fishes on the GBR (Tsai *et al.* 2022) (Fig. S9). However, the model from which eq. S28 and eq. S29 are derived omits variation that is not due to niche differences or species differences in their response to environmental fluctuations (e.g. demographic variance and overdispersion). Nevertheless, comparing our results from eq. S30 and S31 gives us an assessment of how robust the Engen model is to violations of its assumptions in partitioning the variance of relative species abundances.

**Appendix S9: Phylogenetic distance**

An assumption common in community phylogenetics analysis is that closely related species will tend to be more functionally similar than distantly related species, so phylogenetic distance can serve as a proxy for functional diversity and community assembly (Cavender-Bares *et al.* 2009; Connolly *et al.* 2011; Gerhold *et al.* 2015). Here, we wanted to investigate if there is a relationship between species correlations in their responses to environmental fluctuations and species evolutionary history, indicating that closely related species respond similarly to environmental variables. We used the R package *fishtree* (Chang *et al.* 2019) to calculate the phylogenetic distance for 39 out of the 40 species included in our analyses (N.B., there were no phylogenetic data for *Pomacentrus wardi*). Also, *Ctenochaetus striatus* and *Ctenochaetus binotatus* are the two most common *Ctenochaetus* species on the GBR, and we found that their phylogenetic distances with the other species in the study are identical. Therefore, we used the estimates from *Ctenochaetus binotatus* for Ctenochaeuts spp. in our analysis. Then we fitted a simple liner regression model with the package brms (Bürkner 2017).

We found that more closely related species have slightly more positive correlations in their response to environmental fluctuations, with expected correlations decreasing from 0.33 to 0.27 from closely to distantly related species (Fig. S12). However, this relationship is weak (R^2^ = 0.1), suggesting that factors driving environmental correlations are not strongly phylogenetically conserved.

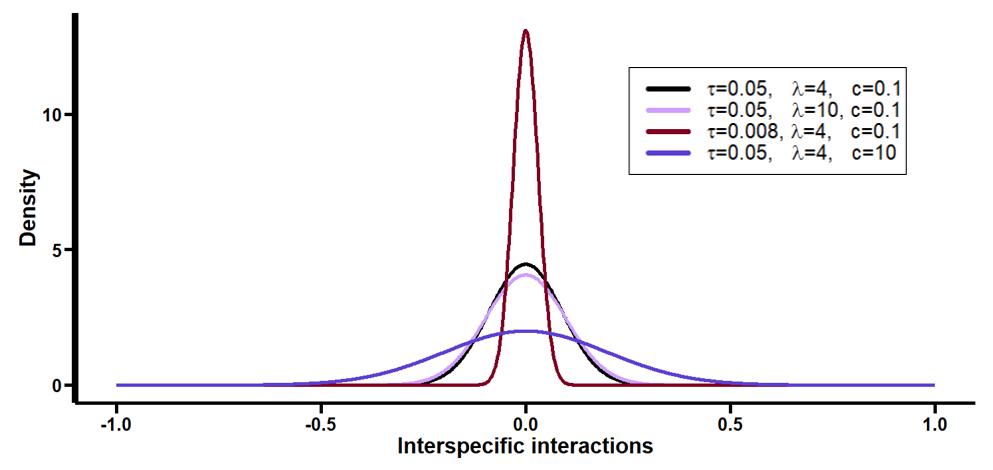
**Figure S1**. Examples of the density of the regularised horseshoe prior for different values of *τ*, *λ* and *c*.

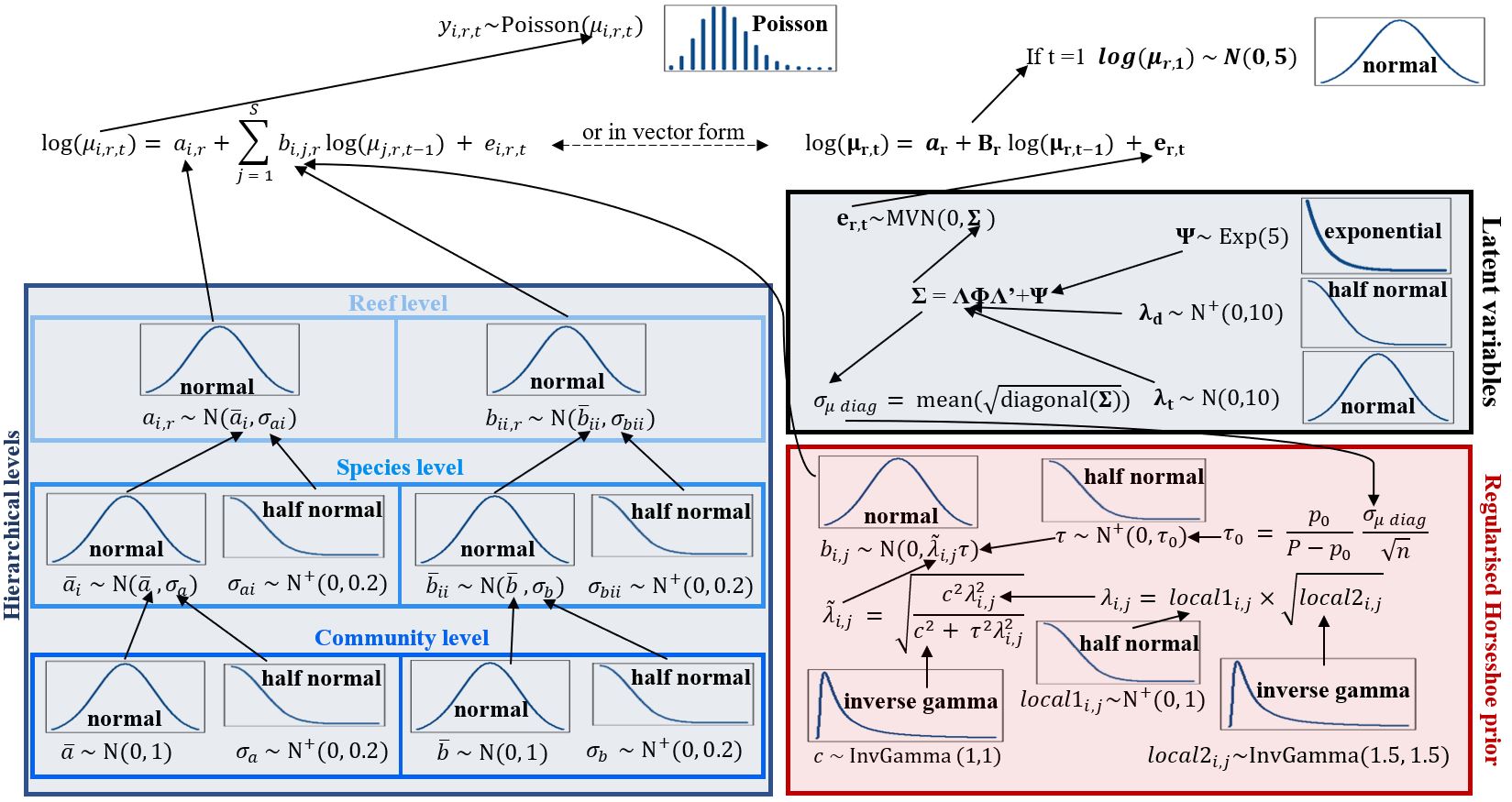
**Figure S2.** Schematic equivalent to Fig. 2 in the main text but here we include the reef random effects as “Reef level” on the Hierarchical levels box (blue box).

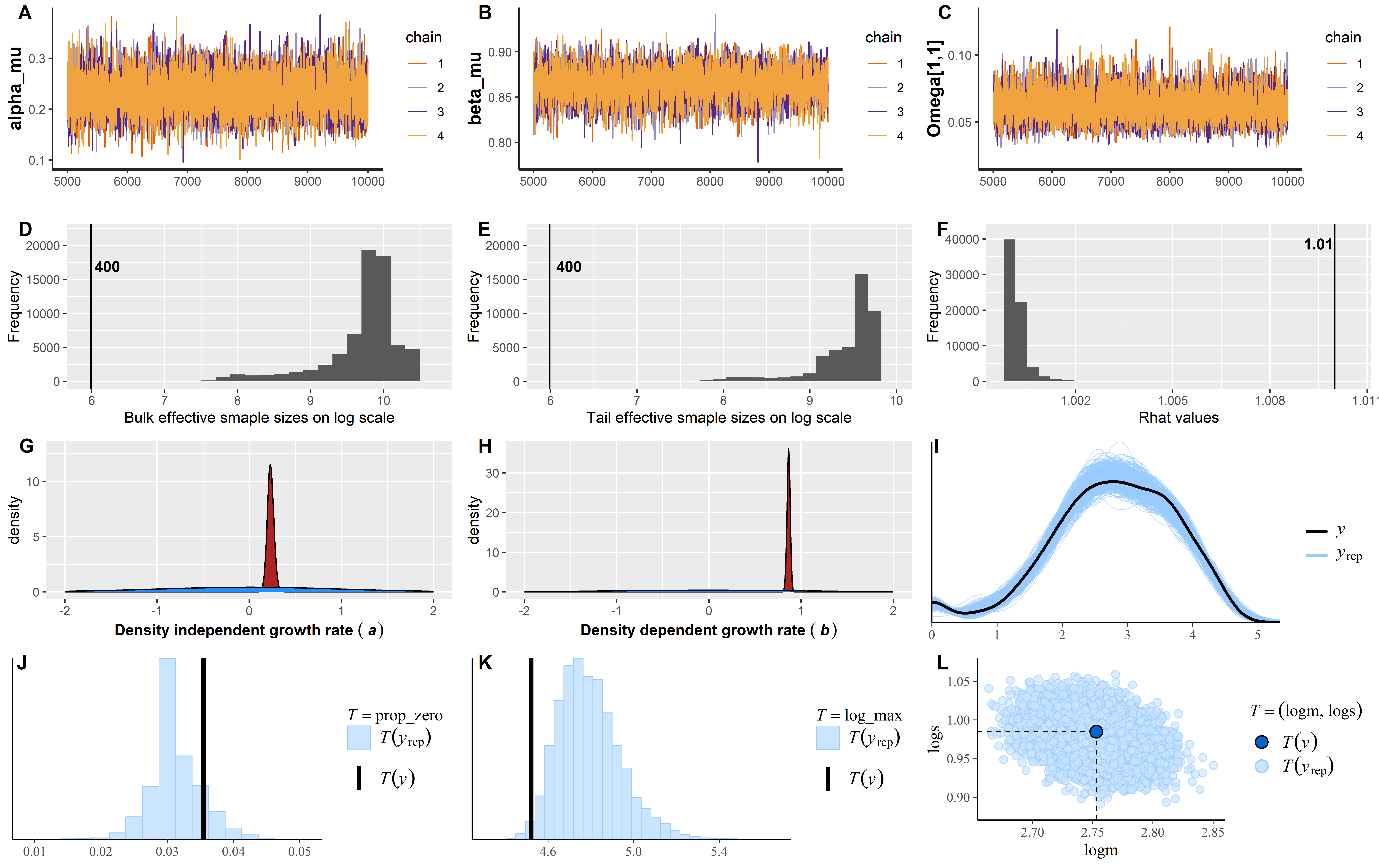

**Figure S3.** Example model diagnostics for our model choice (2 latent variables). **A**, **B** and **C** show the posterior chains for the top level mean intrinsic growth rate (a), the top level mean intraspecific density dependence (b) and the variance for a species in the analysis (Scarus niger), respectively. **D** and **E** show the bulk and tail effective sample sizes (on log scale for easier interpretation), respectively. **F** shows the distribution of R-hats for all the parameters in the model. **G** and **H** show the prior distributions in blue and the posterior distributions in red for the top-level density independent growth rate parameter a and the density dependent growth parameter b, respectively (see Fig. 2). **I** is the posterior predictive check (PPC) for total abundance (NB, the black line represents the log density of the observed data, while the blue lines represent different simulations using the posterior estimates); **J** is the PPC for the proportion of zeros (NB, the vertical black line represents the observed proportion of zeros in the data, while the blue histogram shows the distribution of proportion of zeros for simulated data using the posterior); **K** is the same as J but for the log maximum abundance value; **L** is the PPC for the log mean abundance versus the log standard deviation (NB the dark blue represents the observed value in the data while the light blue the values from simulations using the posterior estimates). **I**-**L** are just examples of PCC for one species (Scarus niger) in the analysis.

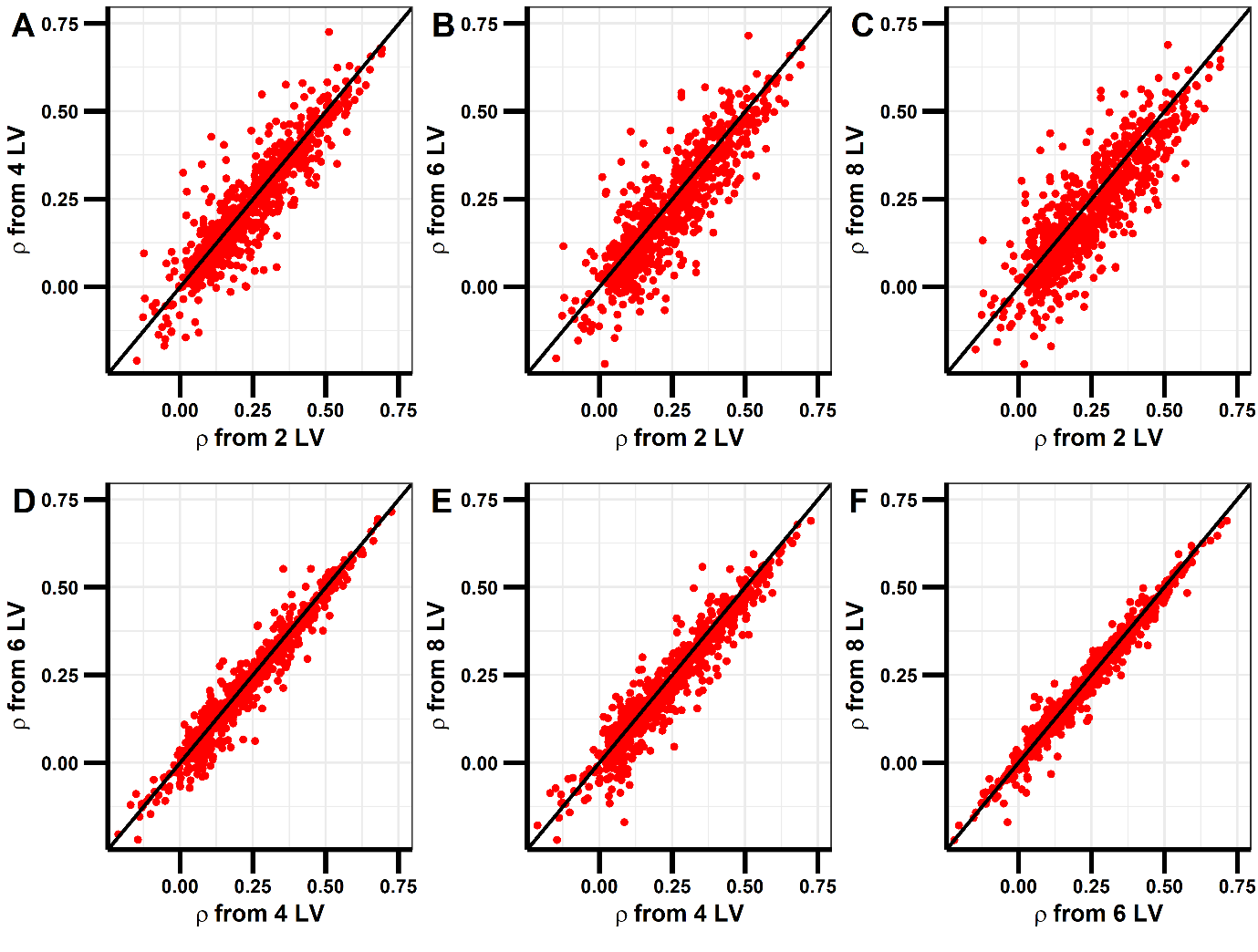

**Figure S4**. Concordance of pairwise environmental correlations for models with different numbers of latent variables. Panel **A** shows species mean pairwise correlations in their responses to environmental fluctuations compared between the model including 2 latent variables versus the model including 4 latent variables. The black line represents the unity line and therefore values close to it indicates that both models estimated the same correlation values. The other panels show the same for the combinations between the model with 2, 4, 6 and 8 latent variables.

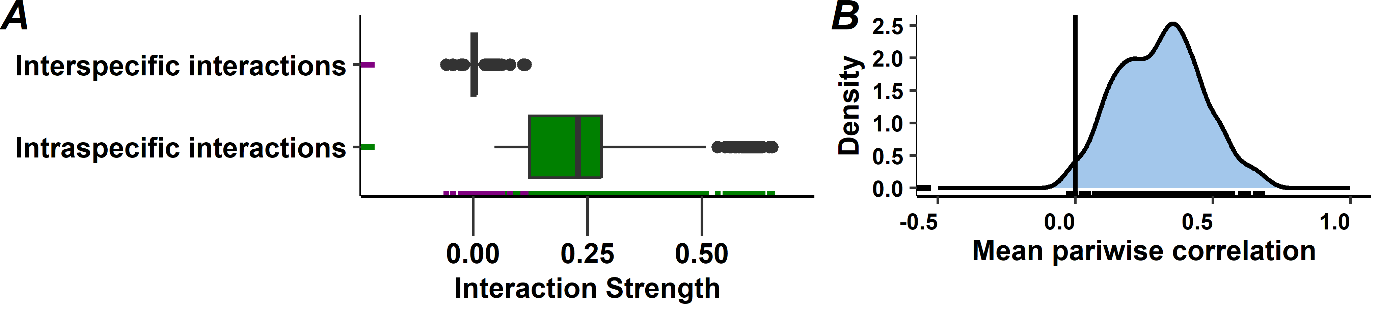

**Figure S5. (A)** Distribution of posterior mean estimates for the elements of the interaction matrix. The diagonal elements are shown in the green boxplot (NB: the diagonal elements are presented here as 1-b_ii_ [i.e., 1 minus the diagonal element], such that zero implies density-independent growth, and positive values imply negative density-dependence). Note that because we modelled reef random effect for the diagonal elements of the interaction matrix, the number of posterior means represented in in the green boxplot are the number of species times the number of reefs (i.e., 41 diagonals, which is 20x41 = 820). The off-diagonal elements are shown in the purple boxplot (NB: b_ij_ = 0 for no interaction; b_ij_ < 0 for negative effects (e.g. competition); b_ij_ > 0 for positive effects (e.g. facilitation)). The marks displayed along the vertical axis represent each of the estimated posterior means for the diagonal elements in green and off diagonal elements in purple. **(B)** Density plot showing the distribution of posterior means of the lower triangular elements of the correlation matrix, calculated from the variance-covariance matrix Σ as $\boldsymbol{P}_{\boldsymbol{ij}}= {\boldsymbol{\Sigma}_{\boldsymbol{ij}}}/{\sqrt{\boldsymbol{\Sigma}_{\boldsymbol{ii}}\boldsymbol{\Sigma}_{\boldsymbol{jj}}}}$. The black marks displayed along the horizontal axis represent each of the estimated posterior correlation means. Note the close resemblance between this figure and Fig. 3B and Fig. 4B in the main text.

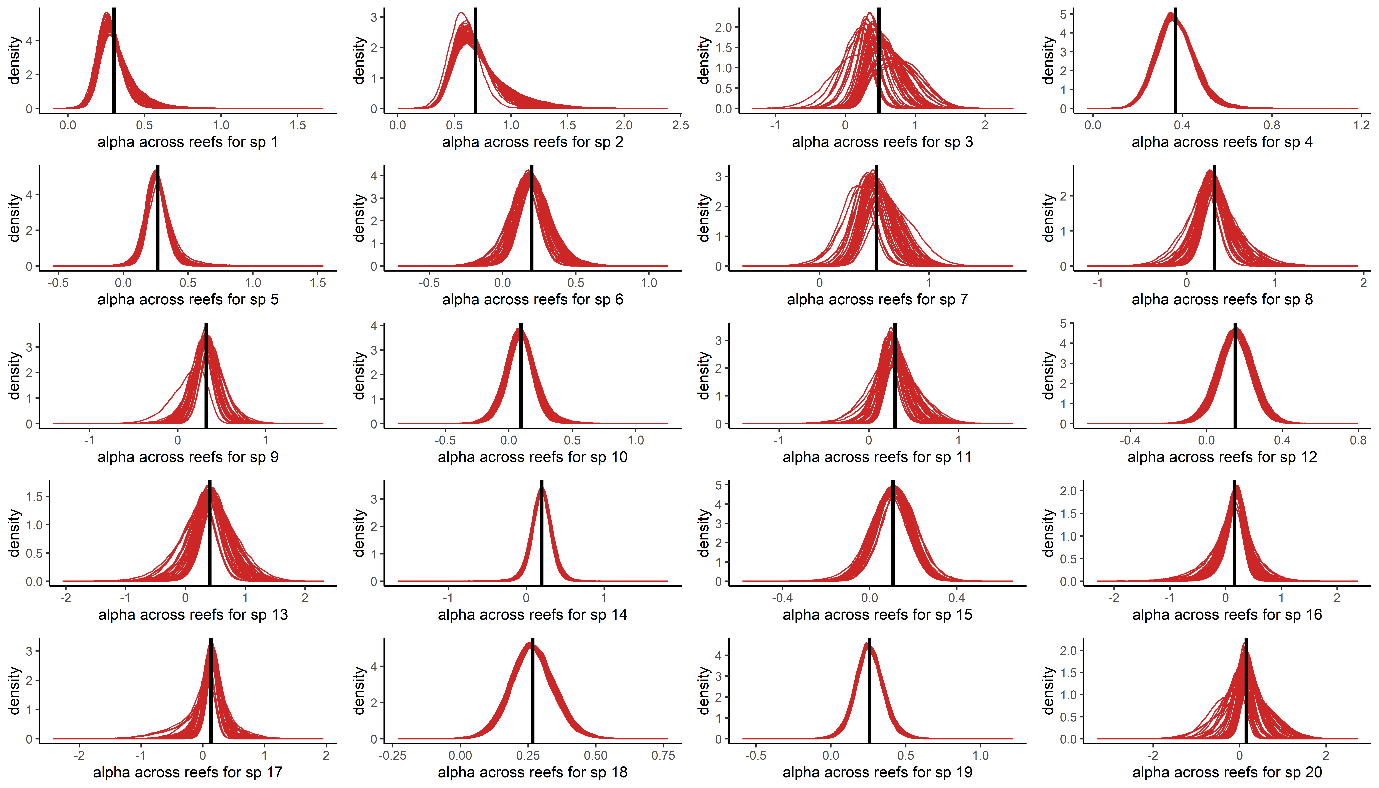

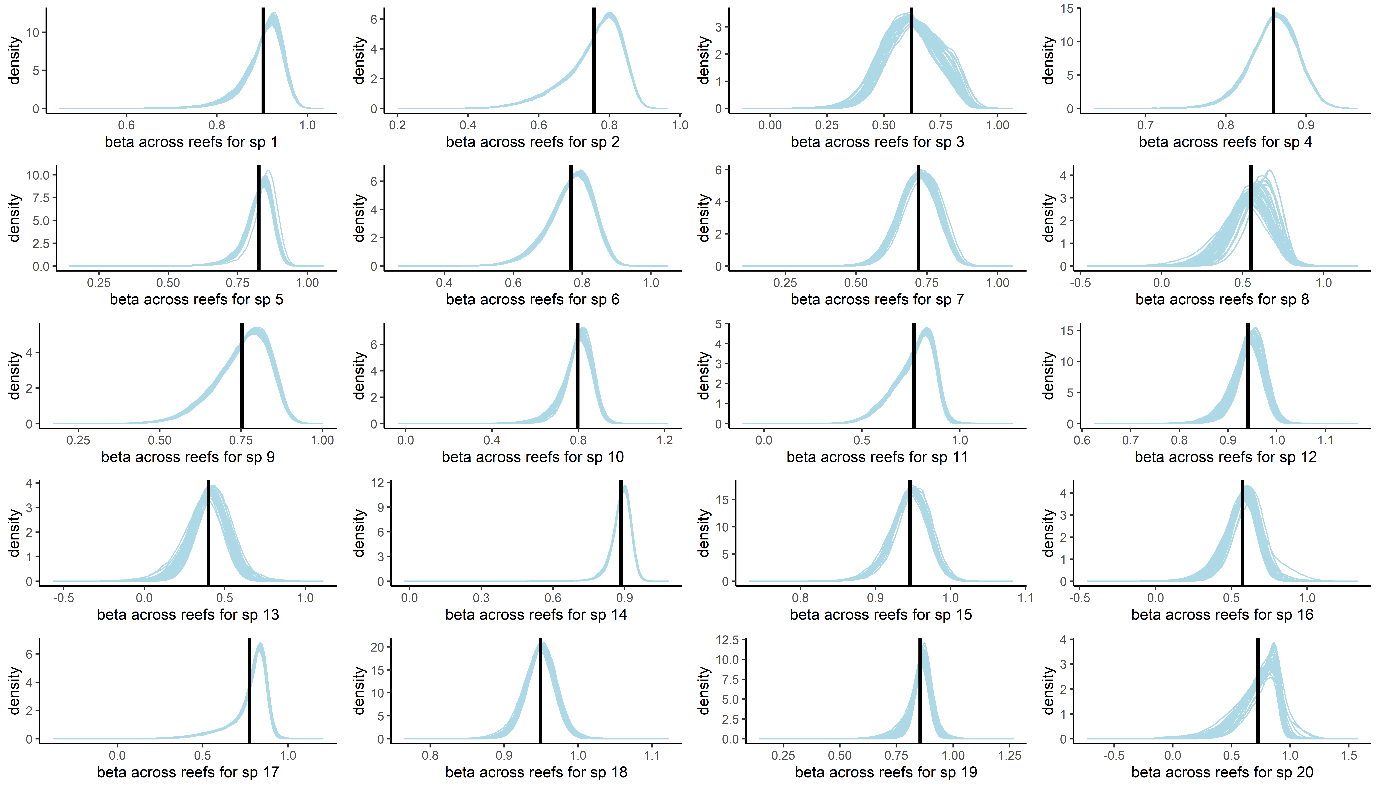

**Figure S6.** Each of the red panels represent a species posterior distribution for the density-independent growth rate parameter alpha across all reefs (41 reefs). Within a panel, one red line represents the posterior distribution of alpha for one reef (i.e., 41 lines). The black vertical line shows the mean alpha at the species level (mean alpha parameter for the level above). The same is represented in the light blue panels for the intraspecific density-dependent growth rate parameter.

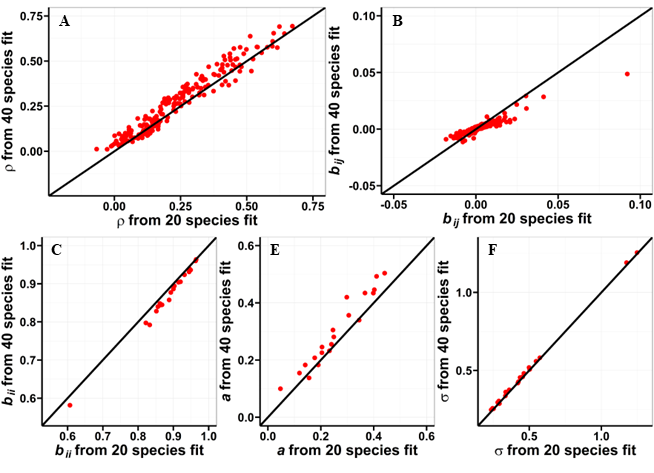

**Figure S7.** Plots comparing the parameter estimates of the model fit to 20 species on the x axis with the same 20 species in the model fit to 40 species on the y axis for the correlation matrix (panel **A**), the interspecific species interactions (panel **B**), the intraspecific density dependence (panel **C**), the intrinsic growth rate (panel **D**) and environmental standard deviation (panel **E**).

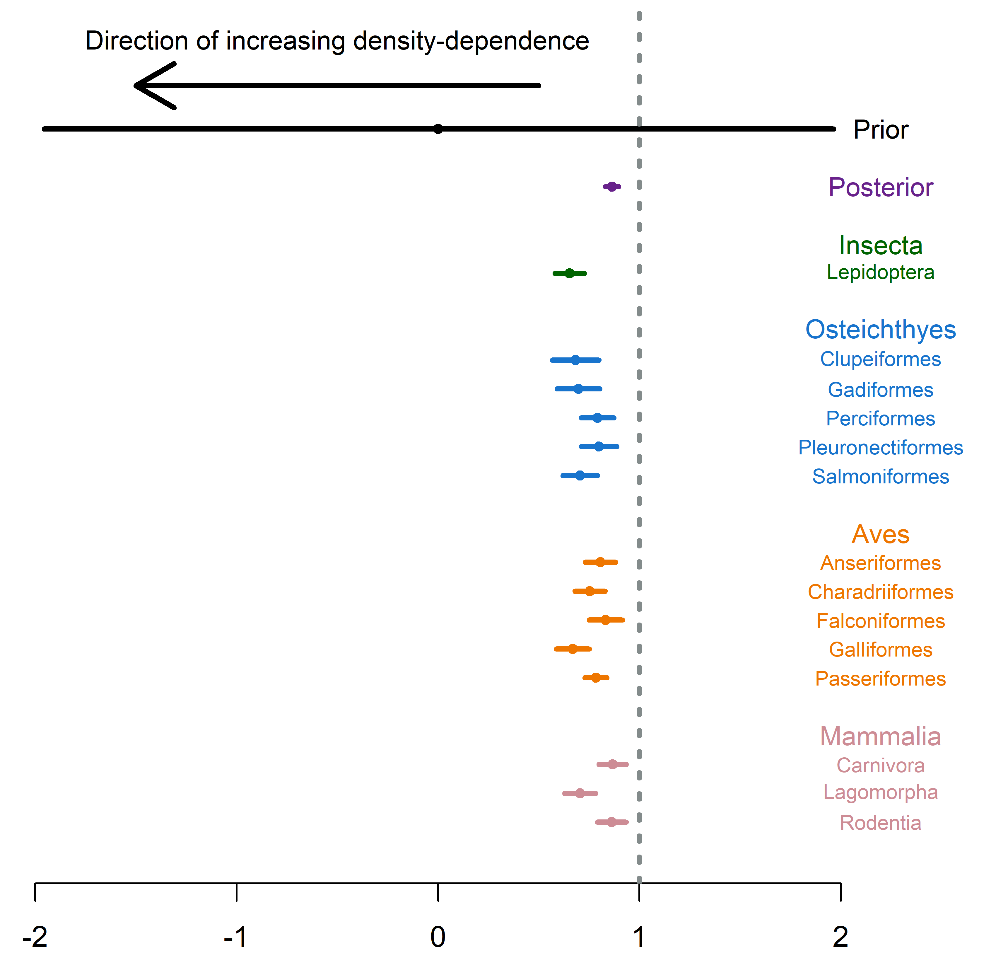

**Figure S8**. Figure adapted from Thibaut & Connolly (2020). The plot shows the mean estimates for intraspecific density dependence. The prior choice at the metacommunity level (see Fig. 2 and S2) with mean (dot) and 95% credible intervals (lines) is represented in black. The same is represented for the posterior estimate at the metacommunity level in purple. The green, blue, orange, and pink dots and line represent the mean and 95% confidence intervals for the intraspecific density dependence estimates calculated by Thibaut and Connolly (2020) for Insect, fishes, birds and mammals.

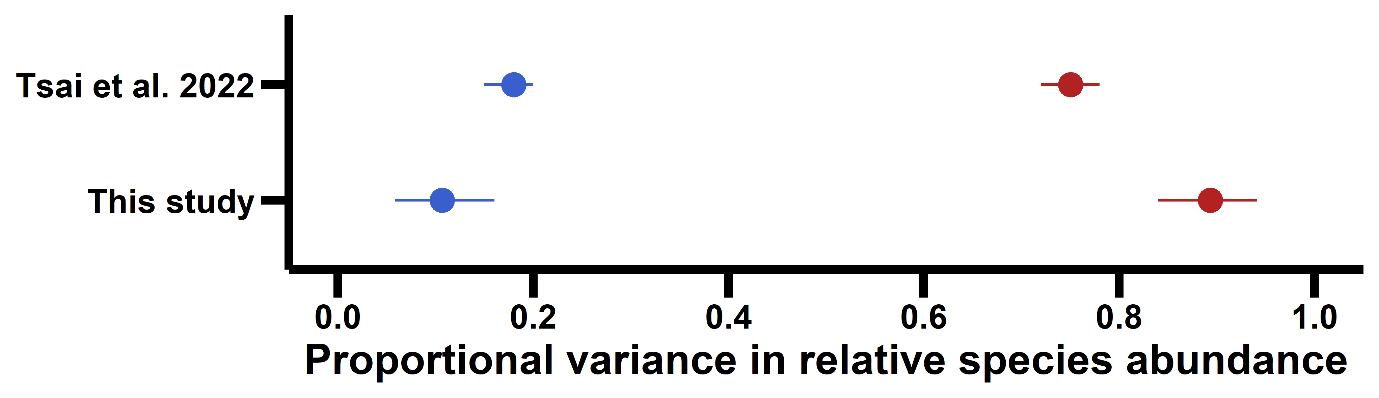

**Figure S9**. Estimated variance components for reef fish on the GBR. For Tsai *et al.* 2022, the dots represent the means and the lines the 95% confidence intervals (frequentist for Tsai *et al.* 2022, Bayesian for this study). Red represents the variance component due to species intrinsic differences. Blue represents the variance component due to environmental fluctuations. In Tsai *et al.* 2022, the variance components were estimated with the VPRSA method. In this study, we used the Gompertz analytical solutions (Ives *et al.* 2003) to estimate the variance due to species differences ($var\left[ {(I-B)}^{-1}A \right]$) and the variance due to environmental fluctuations ($avg\left[ diag\left[ \left( I-B\otimes B \right)^{-1}vec(\Sigma) \right] \right]$).

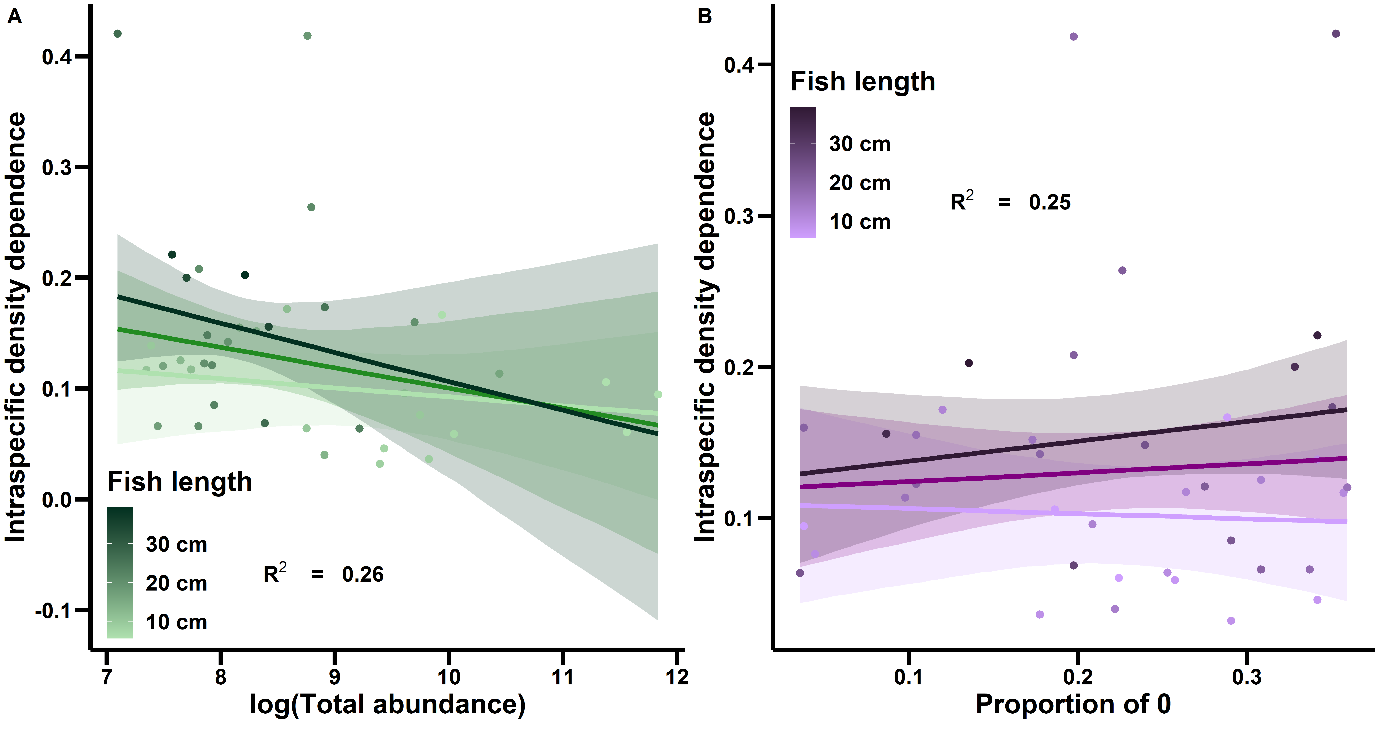

**Figure S10**. Interactive effects of fish length and abundance (measured as total abundance on log scale (**A**) and proportion of zeros (**B**)) on intraspecific density dependence. The points are coloured to represent the gradient of species mean lengths. The light, medium and dark lines represent the model predicted relationship between intraspecific density dependence and the predictor variables (total abundance on log scale in panel **A** and proportion of zeros in panel **B**) for the first (10 cm), median (16.6 cm) and third (21.7 cm) quartile of species mean length, respectively. The corresponding bands represent the 95% credible intervals. The LTMP monitoring program started measuring fish length after 2017. We calculated the mean length for all the measured individuals in each species to obtain a mean length estimate for each species. Even though the estimates of length were from 2017 onwards and our analyses focused on the dynamics between 1995 and 2005, these estimates are specific to the GBR and better characterize the local species than using a theoretical maximum fish length estimate which has been calculated with individuals from other regions (e.g., Fishbase).

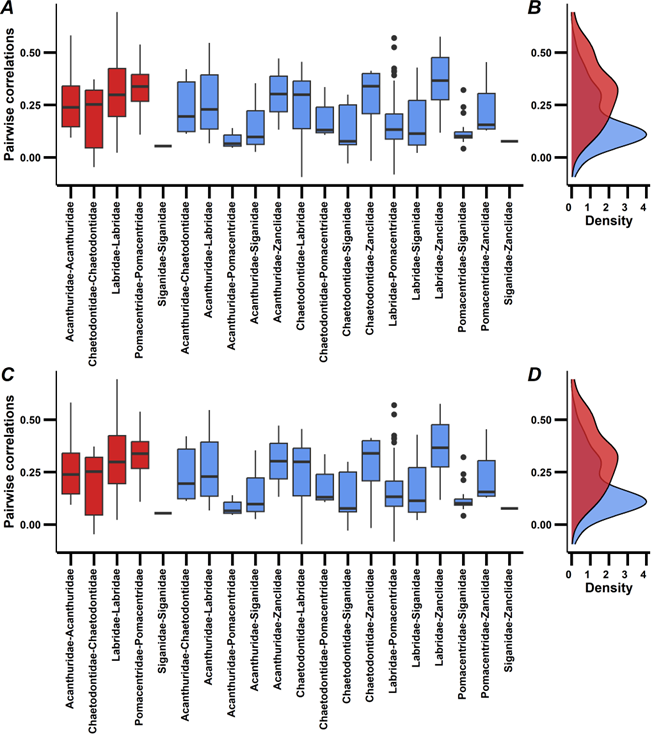

**Figure S11.** In panel **A** the boxplots show pairs of species’ mean correlations in their responses to environmental fluctuations within (in red) and among (in blue) trophic groups. Panel **B** shows the combined correlations within trophic groups and among trophic groups as density distributions in red and blue, respectively. Panel **C** and **D** show the same but at the family level instead of the trophic group level.

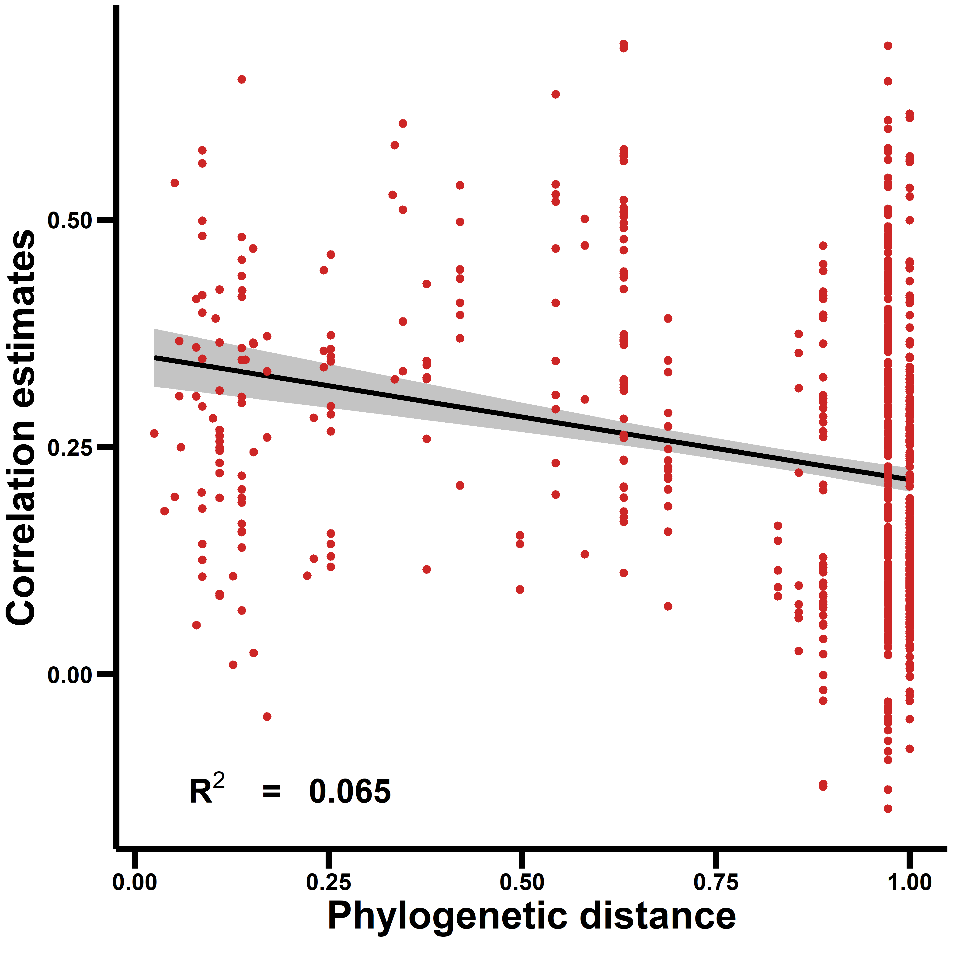

**Figure S12.** Relationship between species mean pairwise correlation estimates and the phylogenetic distance between species pairs. Phylogenetic distances were standardised by dividing all phylogenetic distances values by the maximum distance. Thus, the values are between 0 and 1, with values close to 0 indicating close evolutionary relatedness between species and values close to 1 indicating low relatedness between species. The black line represents a simple linear regression model. The grey band represent the mean 95% credible intervals.

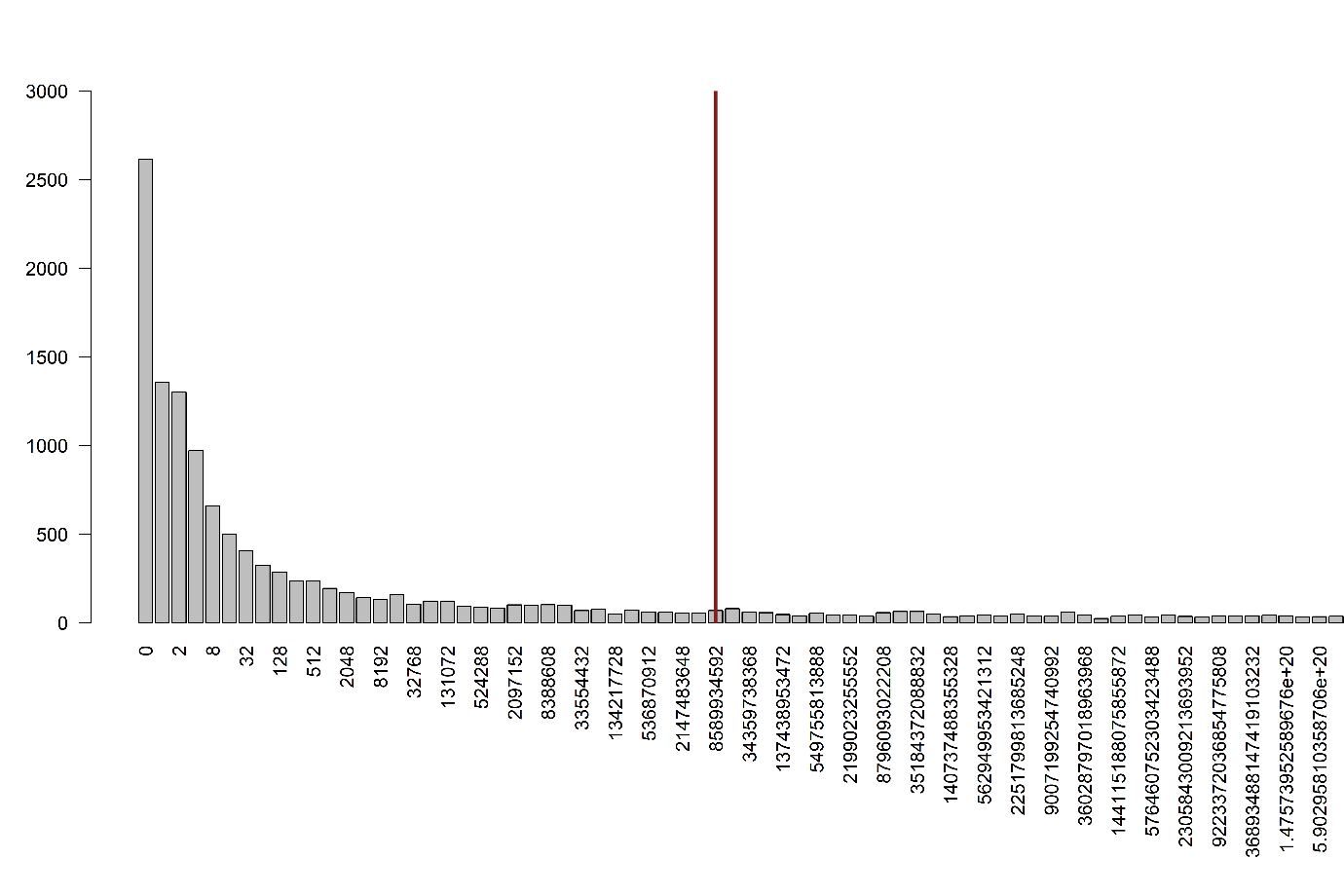

**Figure S13**. Frequency abundance distribution from the simulated data. Each bar represents the frequency for a given abundance for all species across all simulations. Abundances are plotted in log_2_ classes of abundance with the number on the x axis representing the lower limit of an octave (i.e., 0 individuals, 1 individual, 2-3 individuals, 4-7 individuals, etc). The red vertical line represents the approximate number of human individuals on earth for reference.

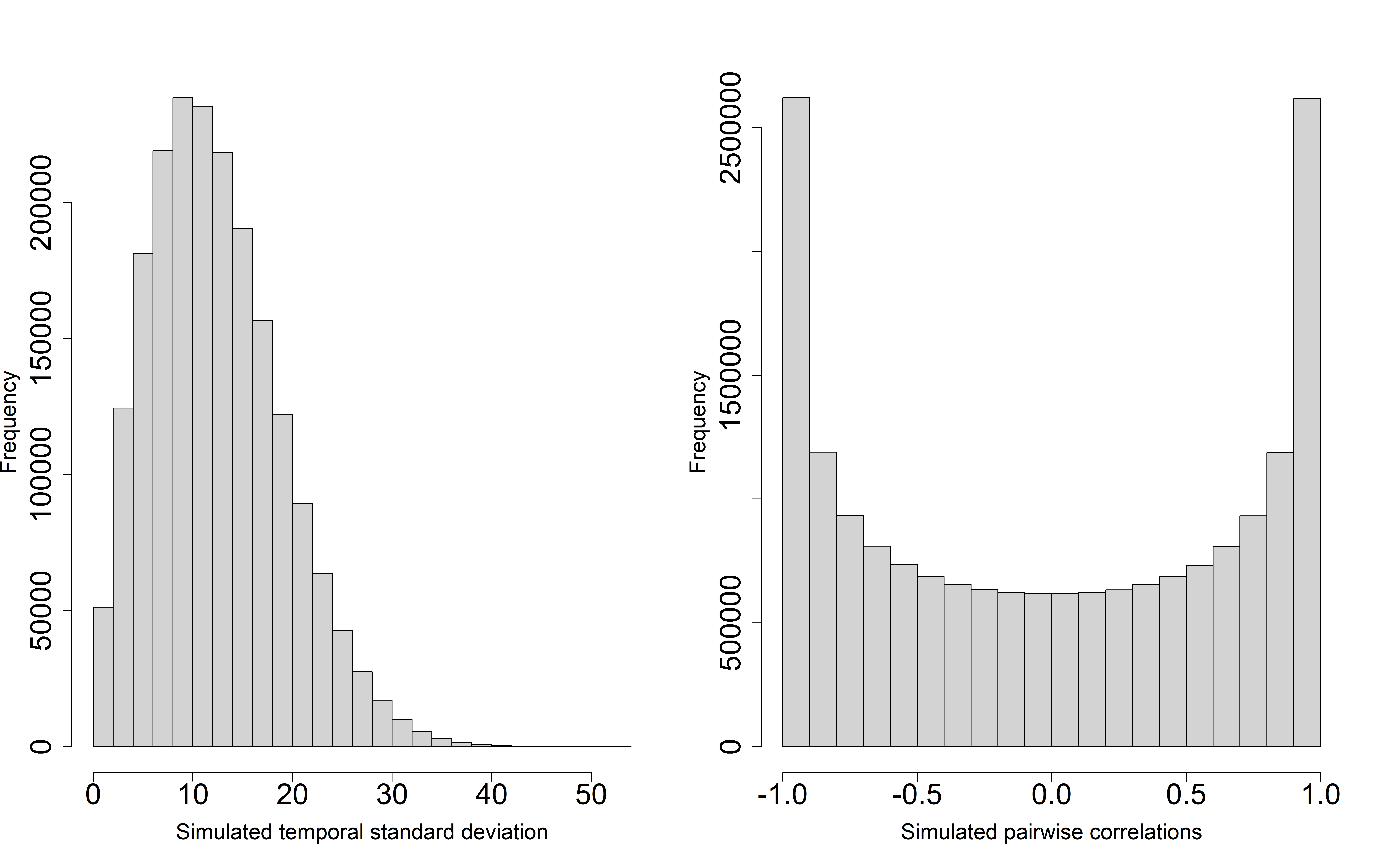

**Figure S14**. The left panel shows the simulated temporal standard deviations (diagonal of the variance-covariance matrix) given our prior choice for the latent variables (see Fig. 2 and S2). The right panel shows the pairwise species correlations in their response to environmental fluctuations given our prior choice for the latent variables (see Fig. 2 and S2).

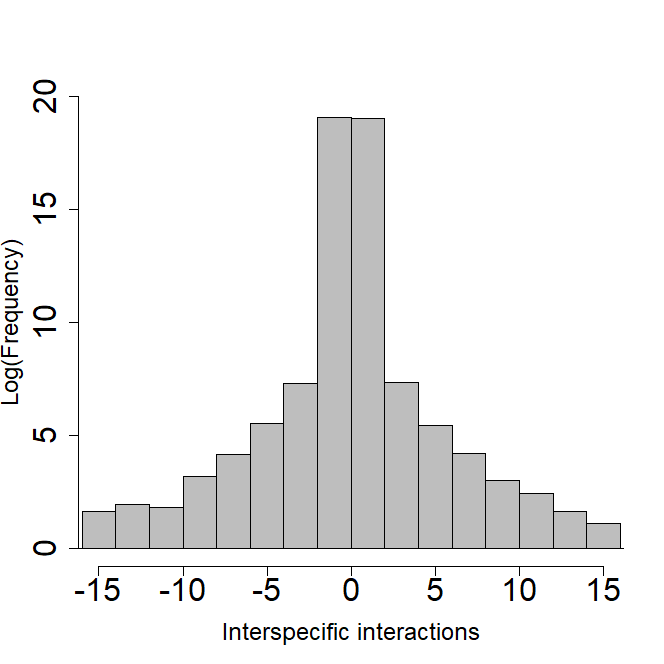

**Figure S15**. The histogram shows the simulated species interspecific interactions (off-diagonal elements of the Interaction matrix **B**, eq. 2) given our prior choice for the regularised horseshoe prior (see Fig. 2 and S2).

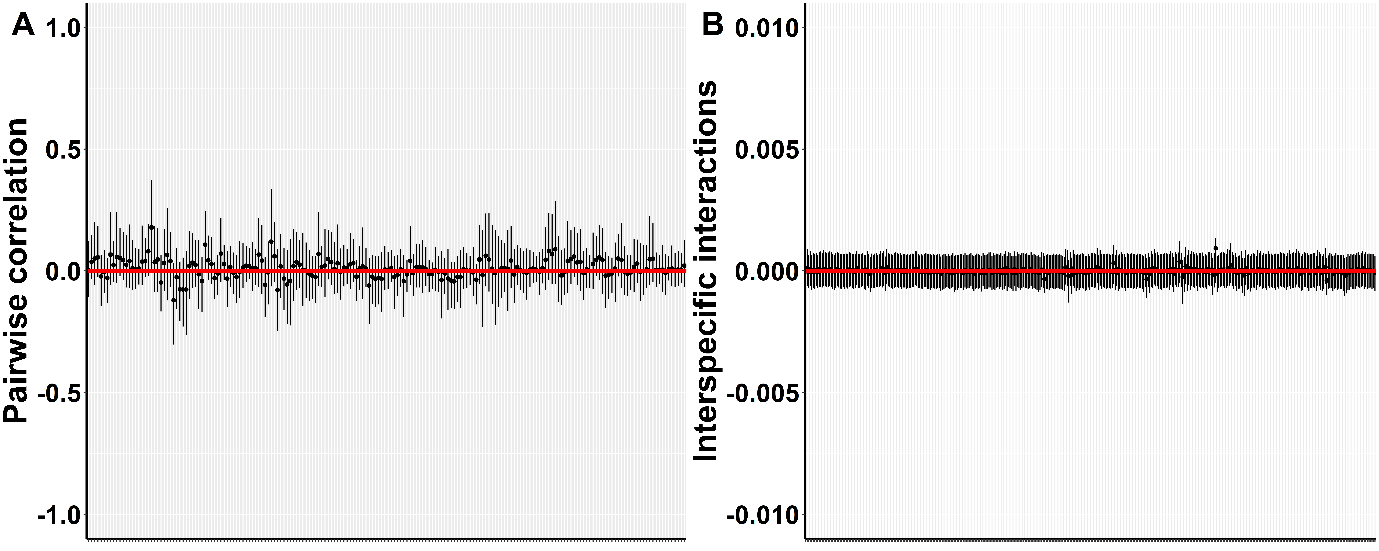
**Figure S16**. Results for *Baseline* case scenario. Panel **A** shows the mean posterior estimates for the species pairwise correlations in their response to environmnetal fluctuarions (black dots), their 95% credible intervals (black lines) and the true values used to simulate the data (red dots). Panel **B** shows the the mean posterior estimates for the species interspecific interactions (black dots) (off-diagonal elements of matrix B in eq. 2), their 95% credible intervals (black lines) and the true values used to simulate the data (red dots).

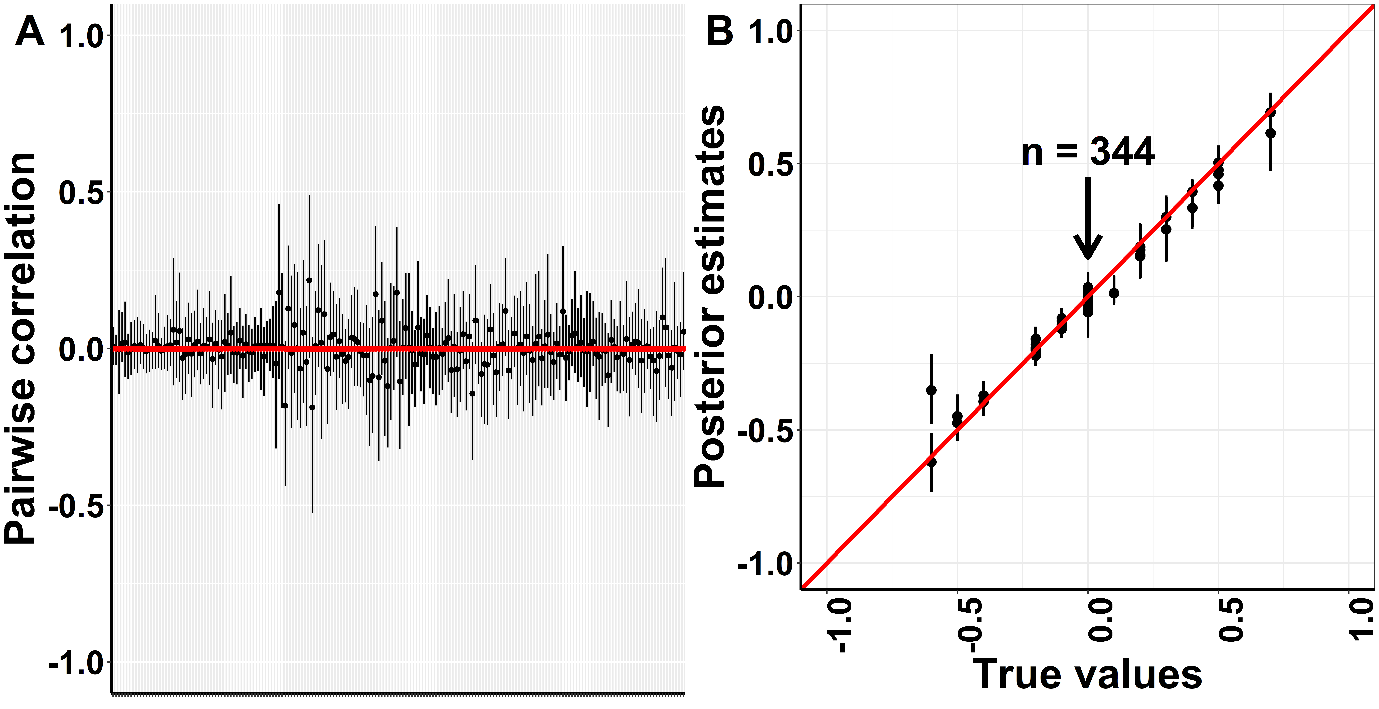

**Figure S17.** Results for Between species interactions case scenario. Panel **A** shows the mean posterior estimates for the species pairwise correlations in their response to environmnetal fluctuarions (black dots), their 95% credible intervals (black lines) and the true values used to simulate the data (red dots). Panel **B** shows the model posterior estimates for the off-diagonal elements of matrix **B** against the true values used to simulate the data for the between species interactions scenario. The black vertical lines represent the 95% credible intervals for the posterior means. The red line represents the unity line. The black arrow points out the clump of zero interactions (n = 344) set to simulate the data.

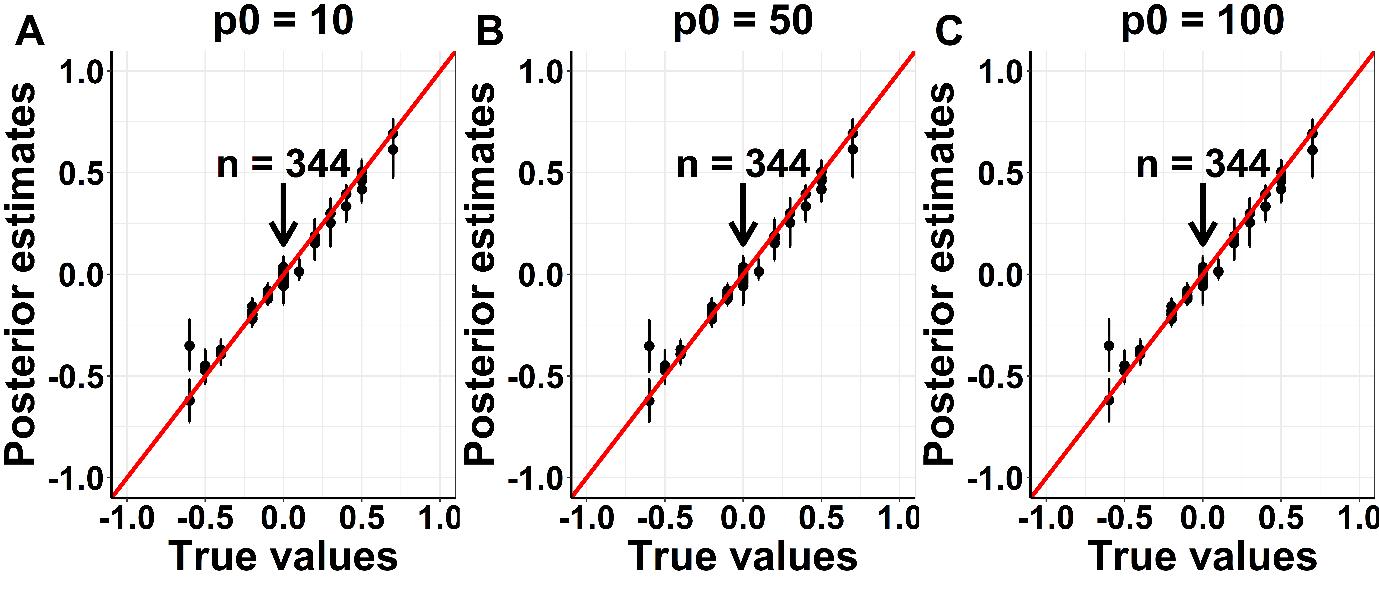

**Figure S18.** Results for Between species interactions case scenario. Posterior estimates for the off-diagonal elements of matrix **B** against the true values used to simulate the data for the between species interactions scenario. The black vertical lines represent the 95% credible intervals for the posterior means. The red line represents the unity line. Each panel represents the model fit with a different prior guess of non-zero parameters, p0. The black arrow points out the clump of zero interactions (n = 344) set to simulate the data and it is the same for the three model fits.

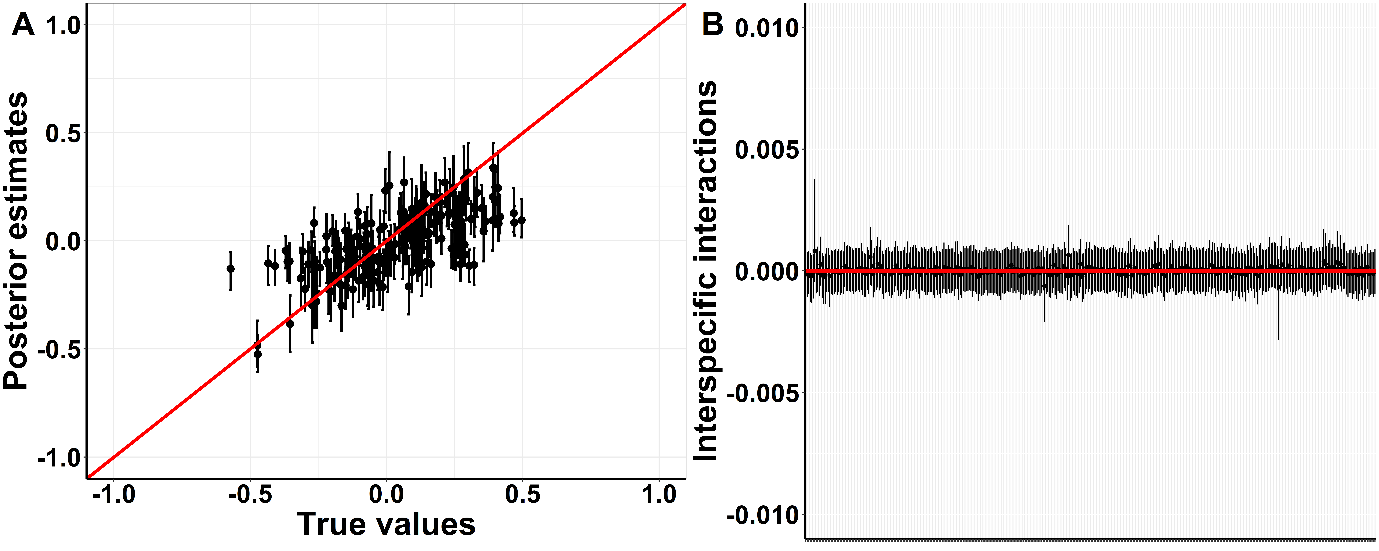

**Figure S19.** Results for Correlated responses to environmental fluctuations case scenario. Panel **A** shows the model posterior mean pairwise correlation estimates for the correlation matrix **P** against the true values used to simulate the data for the correlations scenario. The black vertical lines represent the 95% credible intervals for the posterior means. The red line represent the unity line. Panel **B** shows the the mean posterior estimates for the species interspecific interactions (black dots) (off-diagonal elements of matrix B in eq. 2), their 95% credible intervals (black lines) and the true values used to simulate the data (red dots).

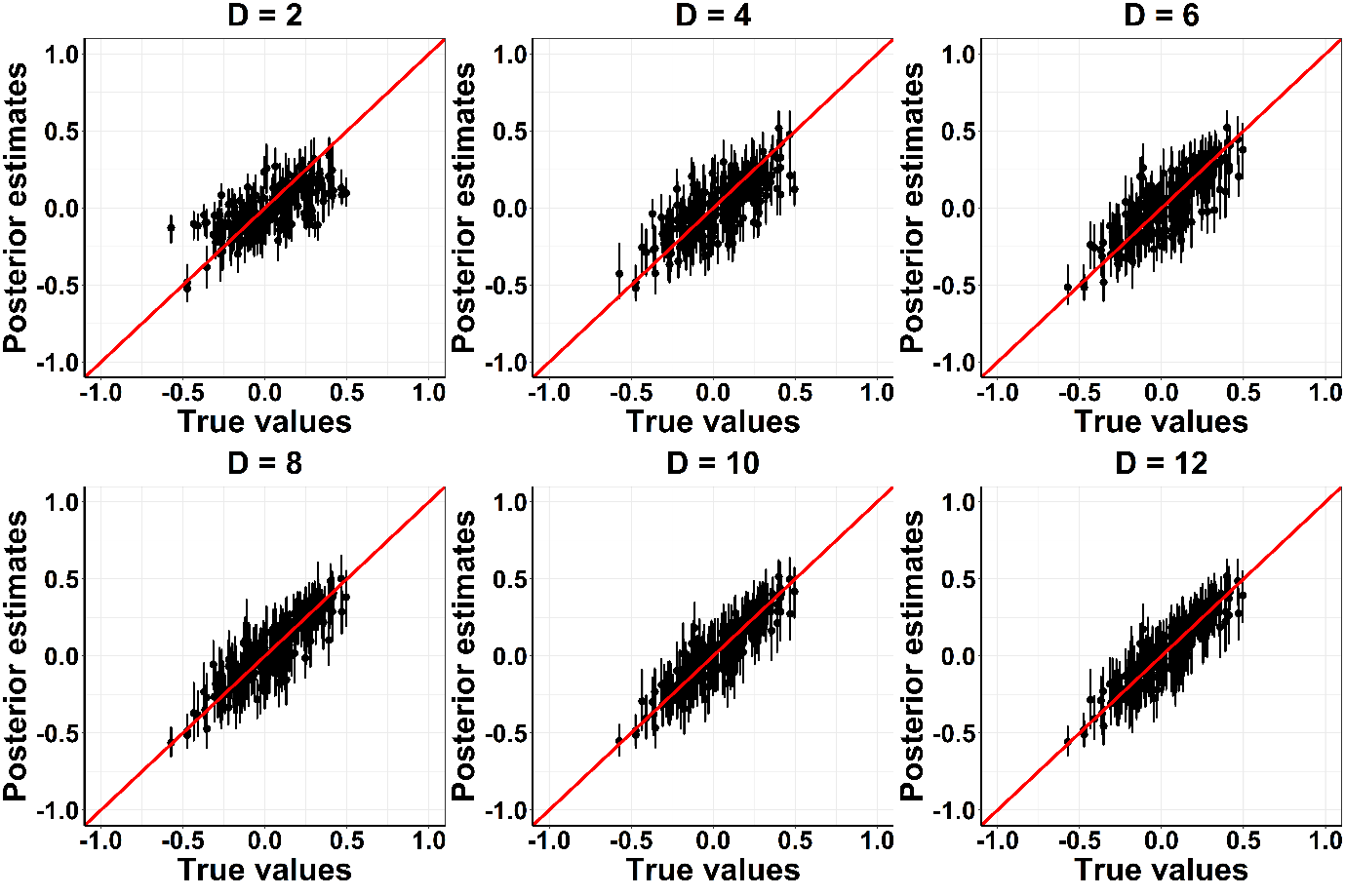

**Figure S20.** Results for Correlated responses to environmental fluctuations case scenario. Posterior mean pairwise correlation estimates for the correlation matrix **P** against the true values used to simulate the data for the correlations scenario. The black vertical lines represent the 95% credible intervals for the posterior means. The red line represent the unity line. Each panels shows the model fit with different number of D latent dimensions.

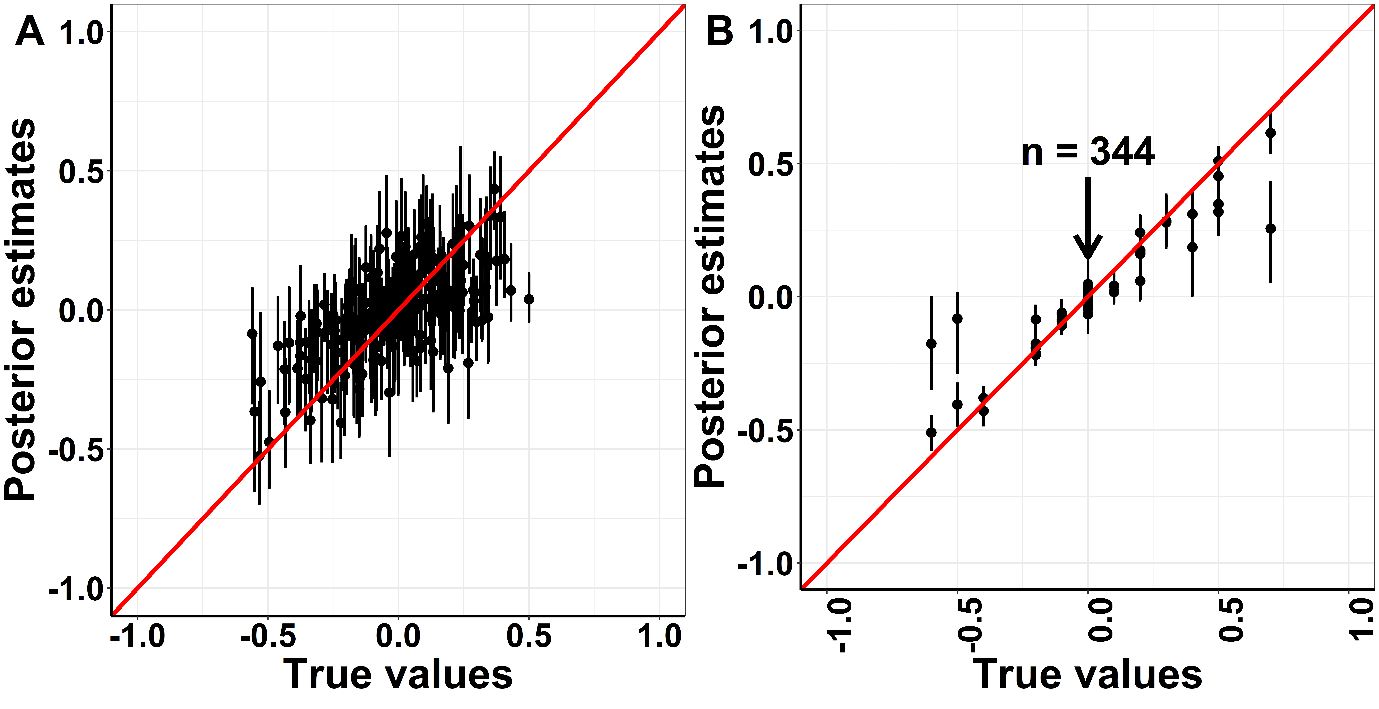

**Figure S21.** Results for Both case scenario.Panel **A** shows the model posterior mean pairwise correlation estimates for the correlation matrix **P** against the true values used to simulate the data for the correlations scenario. The black vertical lines represent the 95% credible intervals for the posterior means. The red line represent the unity line. Panel **B** shows the model posterior estimates for the off-diagonal elements of matrix **B** against the true values used to simulate the data for the between species interactions scenario. The black vertical lines represent the 95% credible intervals for the posterior means. The red line represents the unity line. The black arrow points out the clump of zero interactions (n = 344) set to simulate the data.

**Table S1.** Scientific names and trophic guild of the species analysed in this study. Fish species were placed into trophic groups based on their method of feeding and their impact on the benthos (from published information, e.g., Bellwood & Choat 1990; Froese & Pauly 2006; Green & Bellwood 2009; and field observations)

| Species name | Trophic group |
| --- | --- |
| Scarus niger | Herbivore |
| Chlorurus sordidus | Herbivore |
| Chlorurus microrhinos | Herbivore |
| Ctenochaetus spp. | Detritivore |
| Epibulus insidiator | Invertivore |
| Hemigymnus fasciatus | Invertivore |
| Chaetodon trifasciatus | Corallivore |
| Plectropomus leopardus | Piscivore |
| Gomphosus varius | Invertivore |
| Scarus chameleon | Herbivore |
| Hemigymnus melapterus | Invertivore |
| Scarus frenatus | Herbivore |
| Scarus psittacus | Herbivore |
| Acanthurus nigrofuscus | Herbivore |
| Zebrasoma scopas | Herbivore |
| Scarus globiceps | Herbivore |
| Siganus corallinus | Herbivore |
| Chaetodon rainfordi | Corallivore |
| Chaetodon melannotus | Corallivore |
| Scarus schlegeli | Herbivore |
| Plectroglyphidodon lacrymatus | Herbivore |
| Scarus spinus | Herbivore |
| Chaetodon baronessa | Corallivore |
| Choerodon fasciatus | Invertivore |
| Naso unicornis | Herbivore |
| Halichoeres hortulanus | Invertivore |
| Chaetodon plebeius | Corallivore |
| Scarus altipinnis | Herbivore |
| Scarus rivulatus | Herbivore |
| Siganus punctatus | Herbivore |
| Zanclus cornutus | Invertivore |
| Acanthochromis polyacanthus | Omnivore |
| Amblyglyphidodon curacao | Omnivore |
| Chromis atripectoralis | Planktivore |
| Neoglyphidodon melas | Omnivore |
| Neopomacentrus azysron | Planktivore |
| Pomacentrus lepidogenys | Planktivore |
| Pomacentrus moluccensis | Omnivore |
| Pomacentrus philippinus | Planktivore |
| Pomacentrus wardi | Herbivore |

**Table S2.** Model selection results for models with different number of **D** latent dimensions. Values represent the expected log predictive densities (ELPD), differences in ELPD (elpd_diff) and standard error in ELPD differences (SE_diff) from model selection through leave-one-out cross validation. The preferred model has the value 0 for eldp_diff and SE_diff.

| **Number of latent dimensions** | **elpd_diff** | **SE_diff** | **ELPD** |
| --- | --- | --- | --- |
| **D = 2** | 0.00 | 0.00 | 889.10 |
| **D = 4** | -0.96 | 0.18 | 888.14 |
| **D = 6** | -1.33 | 0.23 | 887.77 |
| **D = 8** | -1.99 | 0.27 | 887.12 |
| **D = 10** | -2.57 | 0.30 | 886.54 |
| **D = 12** | -3.01 | 0.33 | 886.10 |
